# Multi-omics analysis demonstrates a critical role for GLP methyltransferase in transcriptional repression during oogenesis

**DOI:** 10.1101/2021.12.20.473460

**Authors:** Hannah Demond, Courtney W. Hanna, Juan Castillo-Fernandez, Fátima Santos, Evangelia K. Papachristou, Anne Segonds-Pichon, Kamal Kishore, Clive S. D’Santos, Gavin Kelsey

## Abstract

GLP (EHMT1) is a multifunctional protein, best known for its role as an H3K9me1 and H3K9me2 methyltransferase through its reportedly obligatory dimerization with G9A (EHMT2). Here, we investigate the role of GLP in the oocyte in comparison to G9A using oocyte-specific conditional knockout mouse models (*G9a* cKO, *Glp* cKO, *G9a-Glp* cDKO). Loss of GLP in *Glp* cKO and *G9a-Glp* cDKO oocytes re-capitulated meiotic defects observed in the *G9a* cKO; however, there was a significant impairment in oocyte maturation and developmental competence in *Glp* cKO and *G9a-Glp* cDKO oocytes beyond that observed in the *G9a* cKO. Consequently, loss of GLP in oogenesis results upon fertilisation in mid-gestation embryonic lethality. To assess the molecular functions of GLP and G9A, we applied a multi-omics approach, supported by immunofluorescence, to identify changes in epigenomic, transcriptomic and proteomic signatures in cKO oocytes. H3K9me2 was equally depleted in all cKO oocytes, whereas H3K9me1 was decreased only upon loss of GLP. The transcriptome, DNA methylome and proteome were markedly more affected in *G9a-Glp* cDKO than *G9a* cKO oocytes, with transcriptional de-repression associated with increased protein abundance and gains in genic DNA methylation in *G9a-Glp* cDKO oocytes. Together, our findings suggest that GLP contributes to transcriptional repression in the oocyte, independent of G9A, and is critical for oogenesis and oocyte developmental competence.

## Introduction

The mammalian germline is the context for widespread epigenetic changes, in which the somatic epigenetic signature is erased and replaced by a germline signature that is distinct in sperm and oocyte. Prior to the onset of gametogenesis, primordial germ cells undergo extensive erasure of many epigenetic marks, such as DNA methylation (Guibert et al. 2012; Seisenberger et al. 2012) and histone- 3 lysine-9 dimethylation (H3K9me2) (Seki et al. 2005). In the oocyte, epigenetic marks are reset postnatally during oocyte growth, resulting in a unique DNA methylation and histone modification landscape (Hanna et al. 2018a). As the oocyte does not divide during these processes, the oocyte serves as an informative system to study epigenetic regulation. Importantly, resetting of epigenetic marks is essential to support successful oogenesis, the oocyte-to-embryo transition, and subsequent embryonic development (e.g. Kaneda et al. 2004; Andreu-Vieyra et al. 2010; Eymery et al. 2016; Kim et al. 2016; Xu et al. 2019).

G9A (EHMT2) and GLP (EHMT1) are best known as histone methyltransferases, although they modify non-histone targets as well (Rathert et al. 2008). They preferentially function as heterodimers *in vivo* and comprise the main H3K9 mono- and di-methyltransferases in euchromatin (Tachibana et al. 2001; Peters et al. 2003; Rice et al. 2003; Tachibana et al. 2005; Tachibana et al. 2008). Both proteins contain a SET-domain, required for their catalytic activity as methyltransferases and heterodimer formation, as well as an ankyrin repeat domain that enables binding to H3K9me1 and H3K9me2 (Tachibana et al. 2001; Collins et al. 2008). G9A and GLP have been implicated in a number of cellular processes, including gene repression, higher-order chromatin structure, retrotransposon silencing and DNA methylation, not all of which depend on their catalytic activity (Dong et al. 2008; Shinkai and Tachibana 2011; Bittencourt et al. 2014; Auclair et al. 2016; Au Yeung et al. 2019; Jiang et al. 2020). G9A and GLP have been suggested to be inter-dependent, as H3K9me1 and H3K9me2 levels are equally depleted in *G9a* KO and *Glp* KO embryonic stem cells (ESCs) (Tachibana et al. 2005). Moreover, *G9a* and *Glp* mouse knock-outs (KOs) show very similar phenotypes with embryos displaying loss of H3K9me2, growth retardation and embryonic lethality between embryonic day (E)8.5 and E12.5 (Tachibana et al. 2002; Tachibana et al. 2005). As such, studies often do not distinguish between G9A and GLP, or solely consider G9A, resulting in limited knowledge of GLP function. However, recent findings indicate that GLP can act independently of G9A in post fertilisation, revealing a role for GLP in targeting H3K27me2 to the paternal pronucleus in the zygote (Meng et al. 2020).

In the oocyte, loss of G9A impairs maturation and meiosis, with consequences for preimplantation development, leading to partial embryonic lethality (Au Yeung et al. 2019). However, some embryos lacking oocyte-derived G9A develop to term and result in healthy pups. The role of GLP in the oocyte and its impact on embryo development remain unclear. To investigate the importance of GLP in oogenesis and compare its function to G9A, we used oocyte cKO mice for *G9a* and *Glp*, as well as a *G9a-Glp* cDKO. While loss of GLP re-capitulated the meiotic defects observed in the *G9a* cKO (Au Yeung et al. 2019), *Glp* cKO and *G9a-Glp* cKO oocytes showed a significant impairment in oocyte maturation and developmental competence beyond that observed in the *G9a* cKO. Using a multi-omics approach to evaluate changes in H3K9 methylation, gene expression, DNA methylation and protein abundance, we reveal that loss of GLP results in transcriptional de-repression, leading to substantial changes in the oocyte methylome and proteome. Our findings demonstrate that GLP has a unique and essential function in the oocyte.

## Results

### GLP and G9A function during oocyte maturation and meiosis

To analyse the function of GLP in oocytes and compare it to that of G9A, we generated three cKO models. Mice carrying floxed alleles for *G9a* (*G9a* cKO) (Sampath et al. 2007), *Glp* (*Glp* cKO) (Schaefer et al. 2009) or both *G9a* and *Glp* (*G9a-Glp* cDKO) were crossed with a *Zp3*-Cre driver, which is expressed exclusively in growing oocytes after postnatal day 5 (Lan et al. 2004). During oocyte maturation in the ovary, the germinal vesicle (GV) oocyte undergoes a change in chromatin conformation from a non-surrounded nucleolus (NSN) to a surrounded nucleolus (SN) stage that coincides with global transcriptional silencing (Zuccotti et al. 2005). In control animals, immunofluorescence (IF) analysis showed that G9A and GLP are detected throughout the nucleus of NSN oocytes, but were no longer detectable by the mature SN stage (**Supplemental Fig. S1A,B**). In *G9a* cKO oocytes, G9A protein was lost in NSN oocytes, while GLP remained detectable (**Fig. 1A**). In contrast, *Glp* cKO oocytes were depleted for both G9A and GLP, similar to *G9a-Glp* cDKO oocytes (**Fig. 1A**), indicating that GLP is required for G9A stability but not *vice versa*, as was previously shown in mouse ESCs (Tachibana et al. 2005).

**Figure 1.**
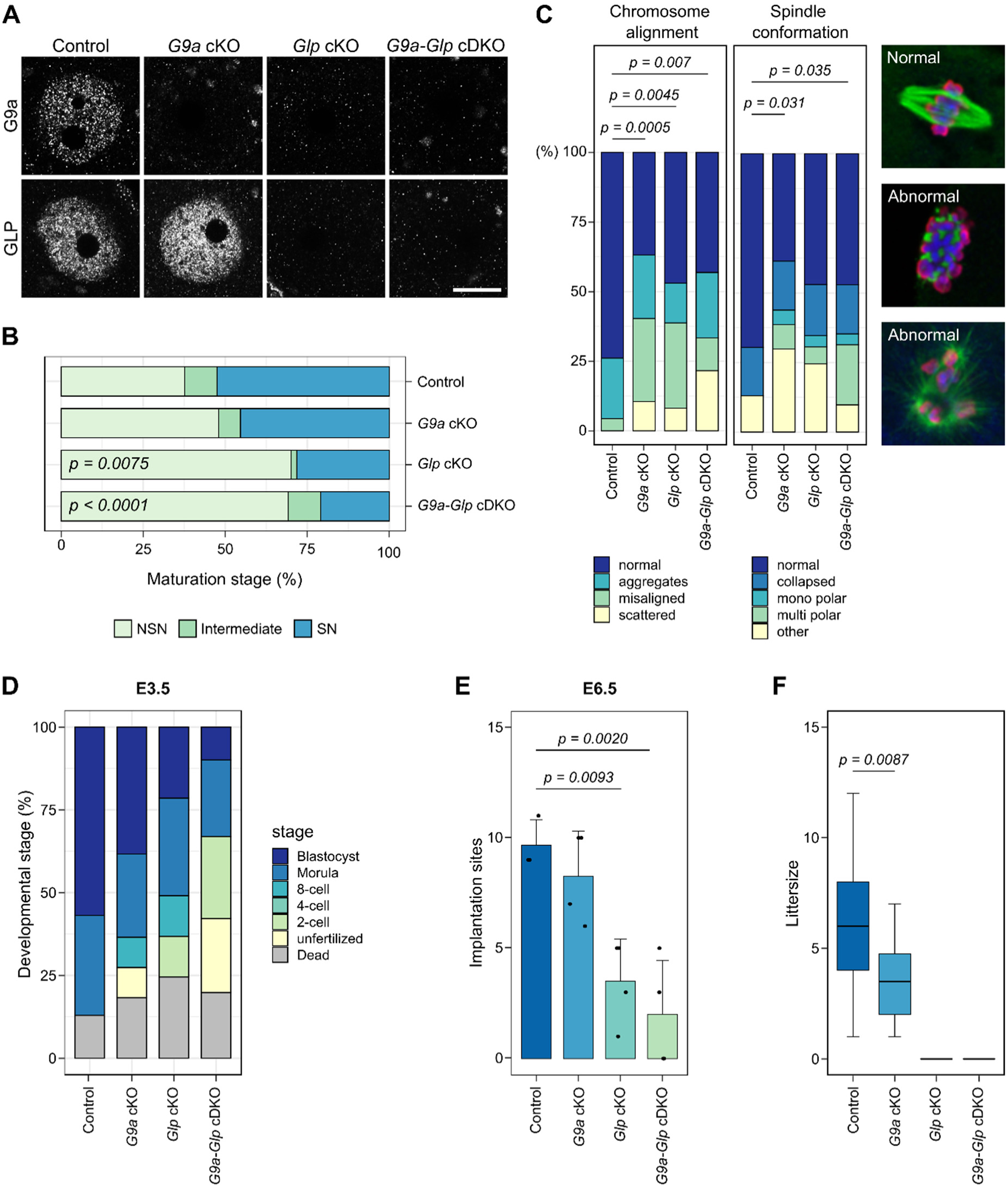
Developmental potential of *G9a* cKO, *Glp* cKO and *G9a-Glp* cDKO oocytes. **A)** Representative images showing IF for G9A and GLP in NSN oocytes. Scale bar: 20 µm. **B)** Stacked bar chart showing percentage of NSN, intermediate and SN oocytes from mice aged 12 weeks. Number of mice/oocytes: Control=5/247, *G9a* cKO=3/160, *Glp* cKO=2/116, *G9a-Glp* cDKO=3/188. Ctrl *vs. G9a* cKO: *P=0.7667*; Ctrl *vs Glp* cKO: *P= 0.0075*; Ctrl *vs G9a-Glp* cDKO: *P<0.0001*; *G9a* cKO *vs G9a-Glp* cDKO *P=0.0058*. **C)** Stacked bar charts showing percentage of chromosome misalignments and spindle abnormalities in MII oocytes. Examples of normal and abnormal spindles are shown in IF images. The spindle is stained with an anti-α-tubulin antibody (green) and the chromatin with DAPI (blue) and anti-pan- histone (red). Number of mice/MII oocytes: Control=5/46, *G9a* cKO=7/57, *Glp* cKO=4/49, *G9a-Glp* cDKO=6/51. **D)** Stacked bar chart showing developmental stage (%) of embryos from cKO females mated with WT males and collected on E3.5. Number of mice/embryos: Control=5/30, *G9a* cKO=5/39, *Glp* cKO=6/35, *G9a-Glp* cDKO=6/29. Blastocysts: Ctrl *vs G9a* cKO *P=0.9939*; Ctrl *vs Glp* cKO P=0.0761, Ctrl *vs G9a-Glp* cDKO *P=0.0073*; One-cell stage: Ctrl *vs G9a-Glp* cDKO *P=0.0406*. **E)** Bar chart showing number of implantation sites scored at E6.5. Dots represent single mice. Ctrl *vs Glp* cKO P=0.0093; Ctrl *vs G9a-Glp* cDKO P=0.0020. **F)** Boxplots showing average litter size of females with cKO oocytes mated with WT males. Number of mice: Control=9, *G9a* cKO=5, *Glp* cKO=3, *G9a-Glp* cDKO=3; Number of litters: Control=60, *G9a* cKO=26, *Glp* cKO=0, *G9a-Glp* cDKO=0; Ctrl *vs G9a* cKO: *P=0.0087*.

Loss of G9A has been shown to affect oocyte maturation and meiosis (Au Yeung et al. 2019). To assess whether loss of GLP has additional effects on developmental capacity of the oocyte, we first analysed the NSN-to-SN maturation rate by staining fully-grown GV oocytes with DAPI and staging them according to chromatin conformation. In *Glp* cKO and *G9a-Glp* cDKO oocytes from 12-week old females the proportion of SN oocytes was significantly lower than in *G9a* cKO and control oocytes (**Fig. 1B**), indicating that loss of GLP has a stronger effect on oocyte maturation than loss of G9A alone.

To determine whether GLP is required for meiosis, we analysed spindle conformation and chromosome alignment of ovulated metaphase II (MII) oocytes after hormonal stimulation. All three cKO models showed an increase in abnormal chromatin configuration and spindle alignment, compared to controls (**Fig. 1C**). Different meiotic abnormalities were observed, ranging from chromosomes that were located together but not aligned (“aggregates”), chromosome alignments where one or several chromosomes were misaligned, to chromosomes scattered throughout the nucleus (**Fig. 1C** and **Supplemental Fig. S1C**). Spindle abnormalities included collapsed, mono- or multipolar spindles. No significant differences were observed between the different cKO models. Taken together, the results show that loss of G9A had mild effects on oocyte maturation compared to the significant impairment caused by loss of GLP, while effects on meiosis were similar.

### Loss of maternal GLP but not G9A results in prenatal developmental arrest

To assess whether loss of GLP also compromises competence of oocytes, we examined developmental progression after fertilisation. Embryos were collected at embryonic day (E)3.5 from cKO females naturally mated with wildtype (WT) C57Bl/6Babr males and scored according to developmental stage. The majority of embryos from control females had reached the blastocyst stage (**Fig. 1D**). In comparison, fewer embryos derived from cKO oocytes progressed to blastocysts, with the proportion of maternal *G9a-Glp* cDKO blastocysts significantly reduced, and the proportion of dead and unfertilized oocytes in maternal *G9a-Glp* cDKO embryos is significantly increased.

To determine the stage of embryo arrest, embryos were collected from superovulated, naturally mated cKO females at E1.5 and cultured *in vitro* for 3 days until day E4.5. Already at E1.5 a difference was observed, all three cKO models having a higher proportion of unfertilized oocytes and 1-cell embryos than controls. The *G9a-Glp* cDKO again showed the strongest phenotype, having significantly fewer 2-cell embryos than controls (**Supplemental Fig. S2A**). Fertilized embryos (1- and 2-cell) were selected for *in vitro* culture: none of the embryos from *Glp* cKO or *G9a-Glp* cDKO oocytes developed to blastocysts, arresting at earlier stages (1 to 4-cell stage) by E3.5 (**Supplemental Fig. S2B,C**). In contrast, 42.9% maternal *G9a* cKO embryos developed at least to morulae by E3.5 (**Supplemental Fig. S2B,C**). However, this was significantly less than controls where 91.1% of embryos reached morula or blastocyst stages by E3.5. These findings reveal that embryos derived from *G9a* cKO oocytes show reduced survival through preimplantation development, whereas preimplantation developmental competence of *Glp* cKO and *G9a-Glp* cDKO oocytes was severely impaired.

A small percentage of embryos from *Glp* cKO oocytes did progress *in vivo* to blastocysts, therefore we examined the implantation and development of embryos after natural mating. In line with findings above, there were significantly fewer implanted embryos at E6.5 from *Glp* cKO and *G9a-Glp* cDKO oocytes than *G9a* cKO and controls (**Fig. 1E**). At E8.5, 3 maternal *Glp* cKO and 6 *G9a-Glp* cDKO embryos were recovered: all were highly abnormal, with no clear tissue types or only extraembryonic tissue (**Supplemental Fig. S2D,E**). In contrast, although some abnormalities (predominantly developmental delay, but also abnormal morphology) were observed among E8.5 embryos from *G9a c*KO oocytes, most appeared normal (**Supplemental Fig. S2D,E**). By E12.5, 55% (11/25) embryos from *G9a* cKO oocytes were grossly abnormal or exhibited developmental delay (**Supplemental Fig. S2F,G**). In *G9a- Glp* cDKO females, with one exception, all embryos had died and only resorption sites were observed (**Supplemental Fig. S2F**).

Finally, we analysed the number of live pups born after mating cKO females with WT males: no pups were born to *Glp* cKO or *G9a-Glp* cDKO females, but *G9a* cKO females did give birth to a mean of 3.46 healthy pups per litter, a significantly reduced litter size compared to controls (**Fig. 1F**). These results confirm previous findings in showing that although loss of G9A in the oocyte affects developmental capacity of pre- and post-implantation embryos, some embryos develop normally resulting in birth of healthy pups (Au Yeung et al. 2019). Our results show loss of GLP in the oocyte severely impairs embryonic development and, although a small proportion of embryos reach the blastocyst stage and can implant, they die *in utero* between E8.5 and E12.5.

### Differential effects of loss of GLP or G9A on H3K9 methylation

GLP and G9A are H3K9 methyltransferases required for establishment of H3K9me1 and H3K9me2 in euchromatin (Tachibana et al. 2001; Tachibana et al. 2002; Tachibana et al. 2005). Therefore, we examined H3K9 methylation in NSN-stage GV oocytes from cKO females (12 weeks) by IF. We tested each antibody in three replicate experiments, each with oocytes from mice from a different litter. In total, we analysed between 13 and 32 NSN oocytes per genotype per antibody (average 22.7). Consistent with previous reports (Peters et al. 2003; Rice et al. 2003), H3K9me1 and H3K9me2 were localized mainly in euchromatic chromatin of NSN oocytes, whereas H3K9me3 was enriched in heterochromatic foci (**Fig. 2A**). Fluorescence intensity of H3K9me1 was significantly reduced in *Glp* cKO and *G9a-Glp* cDKO oocytes compared to controls, but not in *G9a* cKO oocytes; in contrast, H3K9me2 decreased significantly in all three cKOs (**Fig. 2A,B**). A small but significant loss of H3K9me3 was observed in *G9a-Glp* cDKO oocytes (**Fig. 2A,B**). Rather than a direct effect, this may result from loss of H3K9me1, as the H3K9me3 methyltransferase SUV39H requires H3K9me1 as a substrate (Pinheiro et al. 2012). The loss of H3K9me2, but not H3K9me1 and H3K9me3, had previously been shown for *G9a* cKO oocytes using different antibodies, demonstrating the consistency of our IF experiments (Au Yeung et al. 2019)

**Figure 2.**
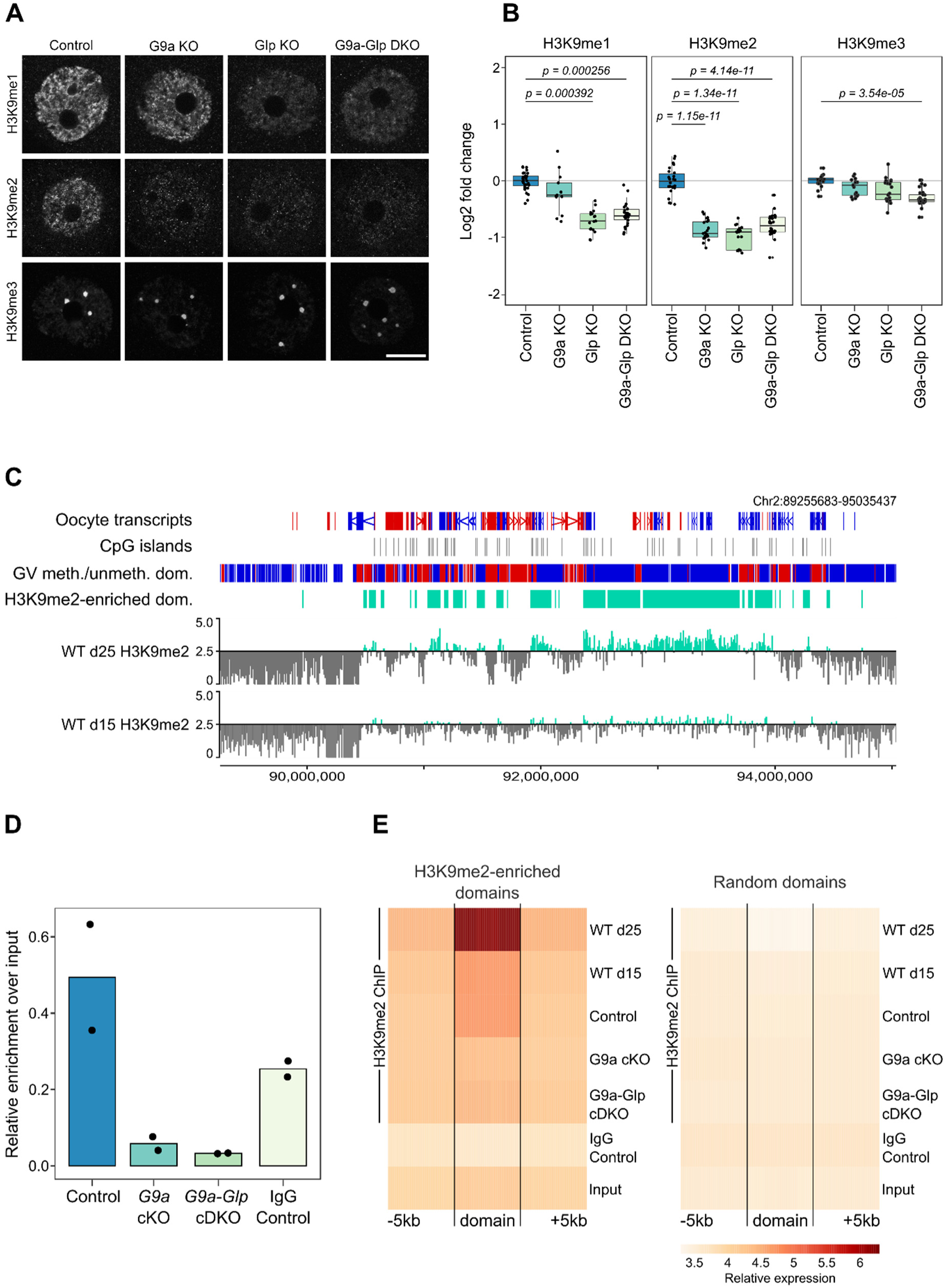
Analysis of H3K9 methylation in *G9a* cKO, *Glp* cKO and *G9a-Glp* cDKO oocytes. **A)** Representative IF images of NSN oocytes for H3K9me1, H3K9me2 and H3K9me3 in control, *G9a* cKO, *Glp* cKO and *G9a-Glp* cDKO oocytes.Scale bar: 20 μm. **B)** Boxplots of quantitation of IF images. Dots represent individual oocytes. Analysis is based on 2-5 mice per genotype and 3-4 replicate experiments for each antibody. **C)** Genome screenshot showing H3K9me2 enrichment for 10kb running windows in WT oocytes from mice aged 15 and 25 days. Y-axis scaling represents log_2_RPKM centred around log_2_RPKM=2.5. Annotation tracks highlight oocyte transcripts, CpG islands, oocyte DNA methylated and unmethylated domains, and H3K9me2-enriched domains (defined as log_2_RPKM>2.5 in d25 GV oocytes). **D)** Boxplots showing relative enrichment over input of H3K9me2 ChIP-seq libraries. Dots represent individual samples. **E)** Probe trend plot showing loss of H3K9me2-enrichment in *G9a* cKO and *G9a-Glp* cDKO oocytes compared to WT and control oocytes in H3K9me2-enriched domains but not random domains.

With H3K9me1 and H3K9me2 affected by loss of G9A and/or GLP, we sought to evaluate the genomic localisation of these marks by ultra low-input native ChIP-seq (ULI-nChIP-seq) in WT GV oocytes from 15 and 25-day old females. We were able to obtain quality ChIP-seq libraries for H3K9me2 (**Fig. 2C**), but not for H3K9me1. By quantifying 10kb running windows, we observed a reproducible H3K9me2 enrichment between replicates from d15 and d25 WT oocytes, with d25 showing greater enrichment than d15 GV oocytes (**Fig. 2C, Supplemental Fig. S3A**). H3K9me2-enriched domains in d25 GV oocytes were defined by merging consecutive 10kb windows with a Log_2_RPKM>2.5 (34,192 or 12.5% of total 10kb windows; 12,514 domains; **Fig. 2C, Supplemental Fig. S3B; Supplemental Table S1**). To link H3K9me2 enrichment to transcription levels, we used published data to define CpG-island (CGI) and non-CGI promoters of low (FPKM<0.1), medium (FPKM 0.1-1.0) and high (FPKM>1.0) expressed genes (Veselovska et al. 2015), and then compared the overlap of these promoters with H3K9me2-enriched or random domains (**Supplemental Fig. S3C**). There was no difference in transcription level between genes localised to H3K9me2-enriched domains or random domains, thus H3K9me2 enrichment does not appear linked to transcriptional repression in oocytes.

For further molecular analysis of GLP function, we used the *G9a-Glp* cDKO model, which had a slightly more severe phenotype than the *Glp* cKO. As this small difference between *Glp* cKO and *G9a-Glp* cDKO oocytes might be caused by residual traces of G9A in the *Glp* cKO oocytes, using *G9a-Glp* cDKO oocytes allowed us to make a clearer distinction between oocytes depleted of both G9A and GLP or only G9A (*G9a* cKO). To assess the loss of H3K9me2 upon depletion of GLP in the oocyte, we generated ChIP-seq libraries of GV oocytes from d25 *G9a* cKO, *G9a-Glp* cDKO and littermate control females. Notably, H3K9me2 libraries from control oocytes were 4.94% of total chromatin, whereas H3K9me2 libraries from *G9a* cKO and *G9a-Glp* cDKO oocytes were only 0.58% and 0.33%, respectively (**Fig. 2D**), confirming that H3K9me2 is almost completely absent from cKO oocytes. Principal component analysis (PCA) of all biological replicates showed that *G9a* cKO and *G9a-Glp* cDKO H3K9me2 replicates clustered apart from control, d15 WT and d25 WT H3K9me2 replicates (**Supplemental Fig. S3D**). When comparing enrichment across H3K9me2-enriched domains (+/-5kb), *G9a* cKO and *G9a-Glp* cDKO showed significant loss of H3K9me2 compared to controls, with enrichment comparable to that in the IgG control and input (**Fig. 2E**). Furthermore, this effect was specific to H3K9me2-enriched domains, as there was no observable difference in enrichment between cKOs and controls across a set of random domains (**Fig. 2E**). There was a similar loss of H3K9me2 in *G9a* cKO and *G9a-Glp* cDKO oocytes, supporting the IF results that G9A is predominantly required for H3K9me2 in oogenesis.

### Dysregulation of proteins associated with meiosis, fertilization and oocyte function in *G9a-Glp* cDKO oocytes

Beside their role as histone methyltransferases, G9A and GLP are known to methylate non-histone proteins and, by doing so, can potentially modulate their function, localisation or stability. To assess whether loss of GLP affects protein abundance in the oocyte, we performed low-input (200 oocytes) quantitative whole-proteome mass spectrometry isobaric labelling analysis of control, *G9a* cKO and *G9a-Glp* cDKO GV oocytes. In total, 21,358 peptides were detected at FDR<1% and 3,182 quantified proteins (**Supplemental Table S2**). Gene ontology (GO) analysis showed enrichment for processes involved in cellular localization and organization, as well as protein folding, metabolic processes and fertilization (**Supplemental Fig. 4A**). Furthermore, among the top 50 most abundant proteins, we detected oocyte-specific proteins such as members of the subcortical maternal complex (PADI6, NLRP5, NLRP14, TLE6, KHDC3, NLRP4F), the zona pellucida (ZP1, ZP2, ZP3), as well as proteins known to be highly abundant in the oocyte (DNMT1, UHRF1).

Comparing *G9a-Glp* cDKO with control oocytes, we identified 187 proteins with a significant change in abundance (*P*<0.05 and Log_2_FC>0.3; **Fig. 3A, Supplemental Table S2**). In contrast, there were only 38 differentially abundant proteins in *G9a* cKO oocytes, of which 21 overlapped with those in *G9a-Glp* cDKO oocytes (**Fig. 3B, Supplemental Fig. 4B**). The majority of changing proteins increased in abundance in both *G9a* cKO (12 down, 26 up) and *G9a-Glp* cDKO oocytes (12 down, 175 up; **Fig. 3A,B**). Although few proteins were identified to have significant changes in abundance in *G9a* cKO oocytes, many proteins changing in *G9a-Glp* cDKO oocytes displayed the same directional trend in *G9a* cKO oocytes (**Fig. 3B**).

**Figure 3.**
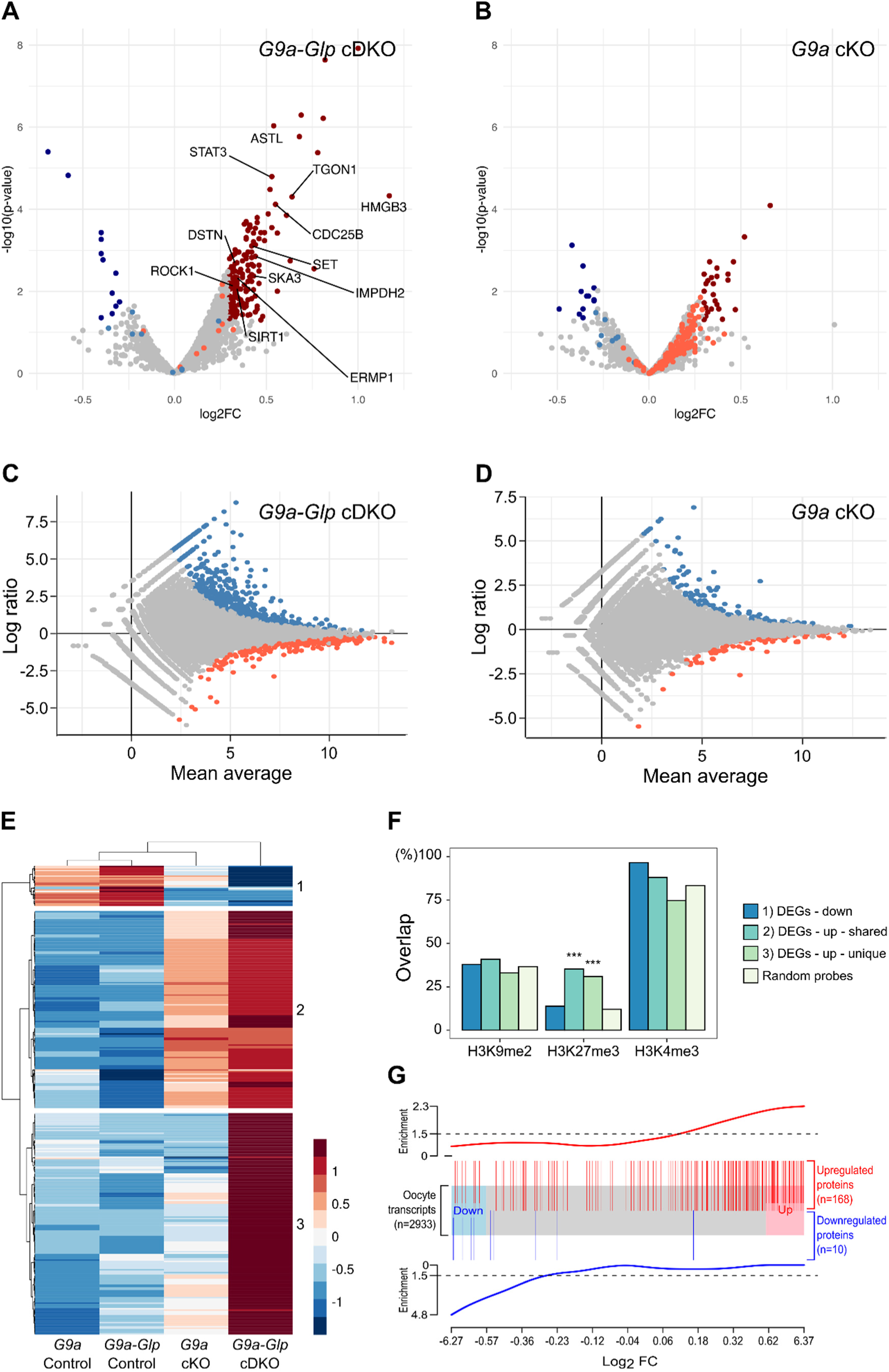
Proteome and transcriptome analysis of *G9a* cKO and *G9a-Glp* cDKO oocytes. **A)** Volcano plot showing differential abundance of proteins in *G9a-Glp* cDKO oocytes compared to controls. Significantly changing proteins (*P*<0.05 and Log_2_FC>0.3) are highlighted in dark blue (decreased abundance) or dark red (increased abundance). Light blue and light red dots indicate proteins changing significantly in *G9a* cKO oocytes. Proteins changing significantly and with oocyte function of interest are labelled. **B)** Volcano plot showing differential abundance of proteins in *G9a* cKO oocytes compared to controls. Significantly changing proteins (*P*<0.05 and Log_2_FC>0.3) are highlighted in dark blue (decreased abundance) or dark red (increased abundance); light blue and light red dots indicate proteins changing significantly in cDKO oocytes. **C)** MA plot showing Log_2_ fold changes in *G9a-Glp* cDKO vs. control oocytes (y-axis) over the mean expression level (x-axis). Differentially expressed genes (DEGs) are highlighted in blue (upregulated) and red (downregulated). **D)** MA plot showing Log_2_ fold changes in *G9a* cKO vs. control oocytes (y-axis) over the mean expression level (x-axis). Differentially expressed genes (DEGs) are highlighted in blue (upregulated) and red (downregulated). **E)** Heatmap showing relative expression levels (RPKM) of up- and down-regulated *G9a-Glp* cDKO DEGs that overlap with *G9a* cKO DEGs. **F)** Barchart showing proportion of DEGs overlapping with histone modifications. DEGs are split according to the clustering analysis of **E**. χ-square analysis comparing DEGs to random probes: P<0.0001***. **G)** Plot showing link between changes in transcript and protein abundance. The x-axis shows Log_2_FC (Control vs. *G9a-Glp* cDKO) of genes present in both transcriptome and proteome datasets (*N*=2933). Blue and red boxes represent down- and up-regulated transcripts, respectively, with a FC>1.5 (Log_2_FC>0.585). Vertical blue and red lines represent differentially abundant proteins (*P*<0.05 and Log_2_FC>0.3; ten proteins not detected in the RNA-seq data are not represented). Enrichment scores, shown above and below, show that downregulated proteins are enriched among downregulated transcripts and upregulated proteins enriched among upregulated transcripts (Spearman correlation *R*=0.43; *P*=2.2x10^-9^).

We detected five known G9A/GLP targets (ACIN1, DNMT1, DNMT3A, MTAL and RUVBL2): none of these were significantly regulated in cKO oocytes, but this is not entirely unexpected as protein methylation does not necessarily influence abundance, but rather may affect activity. GO analysis did not detect any significant enrichment terms amongst the differentially abundant proteins. Even so, we identified several proteins for which change in abundance may be related to the observed oocyte phenotypes. Among these, two were meiotic factors according to the MGI Gene Ontology Browser (CDC25B and SIRT1), while literature research linked a further seven (STAT3, TGON1, SET, IMPDH2, DSTN, SKA3, ROCK1) to meiosis in the oocyte (**Supplemental Table S2**). Furthermore, proteins involved in oocyte maturation (ERMP1) and fertilization (ASTL) showed significantly increased abundance, as well as the candidate oocyte transcriptional regulator HMGB3. Increased abundance of these proteins may underlie the meiotic spindle abnormalities, impaired oocyte maturation and poor fertilization rates observed in *G9a-Glp* cKO oocytes. In line with observations in oocyte and embryo development, the more severe effect in *G9a-Glp* cDKO oocytes suggests unique roles for GLP. The similar but lesser, non-significant changes in protein abundance observed in *G9a* cKO oocytes suggests that loss of G9A can be partially compensated by GLP.

### Transcriptome changes underlie differences in protein abundance in *G9a-Glp* cDKO oocytes

To assess whether the proteome changes may be a consequence of transcriptional changes, we evaluated gene expression by RNA-seq of *G9a-Glp* cDKO, *G9a* cKO and matched control GV oocytes. Differential gene expression was determined by DESeq2 analysis followed by filtering for genes with Log_2_FC>1.5. In line with what was observed in the proteome, the vast majority of differentially expressed genes (DEGs) were upregulated in the *G9a-Glp* cDKO. Again, *G9a-Glp* cDKO oocytes were more severely affected than *G9a* cKO oocytes, with 330 DEGs (301 up, 29 down; **Fig. 3C, Supplemental Table S3**), in contrast to 79 DEGs in *G9a* cKO oocytes (64 up, 15 down; **Fig. 3D**). Of the *G9a* cKO DEGs, 51 overlapped with *G9a-Glp* cDKO DEGs. We identified three clusters of DEGs: 1) downregulated in both *G9a* cKO and *G9a-Glp* cDKO; 2) upregulated in both genotypes; and 3) uniquely upregulated in *G9a-Glp* cDKO (**Fig. 3E**). GO analysis did not show significant category enrichments, but amongst the deregulated transcripts we identified several transcription factors with a known function in the oocyte (*Prmt7, Etv5*), zygotic genome activation (*Zscan4d*) and embryo development (*Klf4, Hoxd1, Lmx1a*; **Supplemental Table S3**). Furthermore, several genes important for oocyte maturation, meiosis and fertilization were deregulated (*Atrx, Fgfr2, Prkcq, Ptgs2, Plac1, Mt1*), which may underlie some of the phenotypic effects we see.

Using the DEG clusters, we assessed the overlap of DEGs with the distribution of histone modifications (**Fig. 3F**). There was no significant enrichment for any of the clusters with H3K9me2, further supporting the conclusion that H3K9me2 does not correlate with transcriptional repression in the oocyte nor explains the differences in severity between the *G9a* cKO and *G9a-Glp* cDKOs (**Fig. 3F**). Upregulated DEGs (clusters 2 and 3) were enriched in H3K27me3, which is localised over untranscribed regions in the oocyte, but not with H3K4me3, which is also broadly localised over untranscribed regions but exclusive with H3K27me3 (Zheng et al. 2016; Hanna et al. 2018). This finding suggests that upregulated DEGs are loci that are transitioning from a repressed to active state in cKO oocytes.

Both our proteome and transcriptome analysis showed a preferential upregulation of protein and transcript abundance in *G9a-Glp* cDKO oocytes. To assess whether changes in transcript expression may be causative for changes in protein abundance, we used gene set enrichment analysis (**Fig. 3G**). Indeed, proteins with increased abundance were enriched for transcriptionally upregulated genes in *G9a-Glp* cDKO oocytes, whereas proteins with decreased abundance were enriched for downregulated genes, although this may not account for all the variation in protein abundance. This indicates that loss of GLP results in deregulated gene expression, which in turn affects abundance of the corresponding proteins. Hence, although only a minority of genes appear to be affected by loss of GLP, these transcriptional changes are likely to be of biological significance.

### Loss of GLP results in local and distinct DNA methylation changes

G9A-mediated H3K9me2 has been linked to DNA methylation (Tachibana et al. 2008; Zeng et al. 2019), although genome-wide analysis only detected local DNA methylation changes in *G9a* cKO oocytes and *G9a* KO embryos (Auclair et al. 2016; Au Yeung et al. 2019). To examine whether loss of GLP affects DNA methylation establishment in the oocyte, we first analysed global DNA methylation (5mC and 5hmC) by IF. We used a previously well-established method (Santos et al. 2013) and performed two replicate experiments with oocytes from different litters (average number of oocytes per genotype = 13). In NSN oocytes, 5mC was observed throughout the nucleus in both euchromatic and heterochromatic regions (**Fig. 4A**); 5hmC was enriched in heterochromatic foci but also present in euchromatin (**Fig. 4A**). No changes in localization of 5mC or 5hmC were observed in cKO oocytes. However, when assessing 5mC fluorescence intensity, there were significant decreases in the *Glp* cKO and *G9a-Glp* cDKO compared to control oocytes, but not for *G9a* cKO oocytes (**Fig. 4B**). No significant changes in 5hmC levels were observed in cKO oocytes, although 5hmC levels also appear reduced in the three cKO models compared to controls. These data suggest that *de novo* DNA methylation may be impaired upon loss of GLP but not G9A in oocytes.

**Figure 4.**
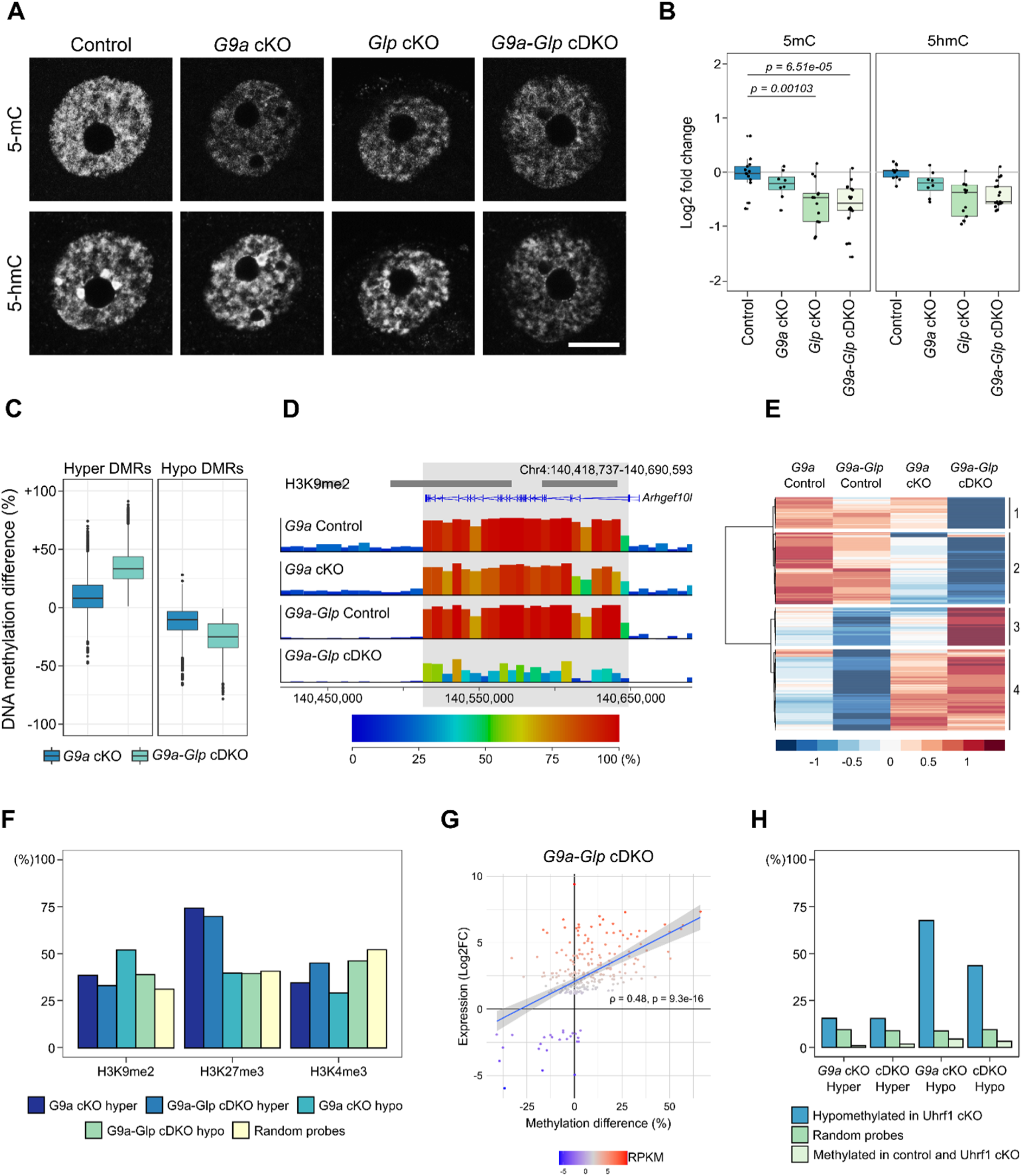
DNA methylation changes in *G9a* cKO and *G9a-Glp* cDKO oocytes. **A)** Representative IF images of NSN oocytes showing 5mC and 5hmC in Control, *G9a* cKO, *Glp* cKO and *G9a-Glp* cDKO oocytes. **B)** Boxplots of quantitation of IF images. Dots represent individual oocytes. Analysis is based on 2 mice per genotype and 2 replicate experiments for each antibody. **C)** Boxplot showing the methylation difference of *G9a-Glp* cDKO DMRs (hyper and hypo) in controls vs *G9a* cKO and controls vs *G9a-Glp* cDKO oocytes. **D)** Genome screenshot showing example of a region uniquely hypomethylated in *G9a-Glp* cDKO oocytes spanning the *Arhgef10l* transcript. Annotation tracks show position of H3K9me2-enriched domains and oocyte transcripts. **E)** Heatmap showing clustering analysis of *G9a-Glp* cDKO hypo- and hypermethylated domains. **F)** Barchart showing percentage overlap of DMRs and random probes with H3K9me2, H3K37me3 and H3K4me3. H3K9me2: *G9a* cKO hyper *adj. P*=0.0136, *G9a* cKO hypo *P<0.0001, G9a-Glp* cDKO hypo *P<0.0001*; H3K27me3: *G9a* cKO hyper *P<0.0001, G9a-Glp* cDKO hyper *P<0.0001*; H3K4me3: all genotypes *P<0.0001*. **G)** Plot showing correlation between methylation changes (% methylation difference) and expression changes (Log_2_FC) of DEGs in *G9a-Glp* cDKO oocytes. Relative expression levels (RPKM) of each gene are indicated by the colour scale. Spearman correlation is shown. **H)** Barchart showing percentage overlap of DMRs with 100-CpG windows that are hypomethylated in *Uhrf1* cKO oocytes (>20% methylation difference, (Maenohara et al. 2017)), random 100-CpG windows and 100-CpG windows that are highly methylated (>75%) in WT and *Uhrf1* cKO oocytes. *G9a* cKO hyper *adj. P=0.0404*, *G9a-Glp* cDKO hyper, *G9a* cKO hypo and *G9a-Glp* cDKO hyper *P<0.0001*.

We then explored changes in DNA methylation in greater resolution by whole-genome bisulphite sequencing (BS-seq) of control, *G9a* cKO and *G9a-Glp* cDKO GV oocytes. In contrast to decreased 5mC seen by IF, BS-seq did not show significant global changes in DNA methylation (**Supplemental Fig. S5A**). An explanation for this apparent discrepancy is that the genomic regions assessed by the two methods differ: IF signal reports both euchromatic and heterochromatic fractions of the genome, while BS-seq data is enriched in euchromatic and non-repetitive regions that can be uniquely mapped and analysed. These differences highlight the value of using both methods to understand genome-wide patterns of DNA methylation and may explain previous contradictory findings (Au Yeung et al. 2019; Zeng et al. 2019).

Although global DNA methylation levels as detected by BS-seq did not differ, local changes in DNA methylation were observed after binning the genome into consecutive windows of 100 CpG sites (100- CpG windows) for analysis. Differential methylation analysis identified 9,187 DMRs in *G9a-Glp* cDKO oocytes (4.88% of all analysed 100-CpG windows), of which 4,184 (45.5%) were hypermethylated (mean methylation difference: 35.1%) and 5,003 (54.5%) hypomethylated (mean methylation difference 25.6%) (**Supplemental Fig. S5B; Supplemental Table S4**). Methylation was more affected in *G9a-Glp* cDKO than in *G9a* cKO oocytes, with >7 times more DMRs identified: there were 1,252 DMRs (0.67% of analysed 100-CpG windows) in the *G9a* cKO, of which 432 (34.5%) were hypermethylated (mean methylation difference: 39.4%) and 820 (65.5%) hypomethylated (mean methylation difference: 33.8%) (**Supplemental Fig. S5C**). The majority of *G9a* cKO DMRs overlapped *G9a-Glp* cDKO DMRs (**Supplemental Fig. S5D**). Furthermore, not only did the *G9a-Glp* cDKO have more DMRs, the methylation difference of these DMRs was greater in *G9a-Glp* cDKO oocytes (**Fig. 4C**). This indicates that while both G9A and GLP are required for normal DNA methylation establishment in the oocyte, GLP can partially compensate for loss of G9A, but it does not exclude the possibility that GLP has an additional G9A-independent role in DNA methylation.

Between 21 and 38% of 100-CpG windows identified as DMRs cluster and form larger regions that, in both genotypes, can span genes (**Fig. 4D**, **Supplemental Fig. S6** and **Supplemental Table S4)**. In total, there were 707 hypo- and 869 hyper-methylated domains in *G9a-Glp* cDKO oocytes, with fewer in the *G9a* cKO (140 hypo- and 66 hyper-methylated domains). To see whether the additional changes in the *G9a-Glp* cDKO are unique or whether similar (non-significant) trends are present in the *G9a* cKO, we performed unsupervised cluster analysis of DNA methylation of the cDKO domains (**Fig. 4E**). For both hypo- and hypermethylated domains, we identified domains common to both cKOs (523 hypo, 590 hyper) and domains unique to the *G9a-Glp* cDKO (229 hypo, 279 hyper; **Fig. 4D,E; Supplemental Fig. S6**). This indicates that the majority of effects seen in the *G9a-Glp* cDKO are also present in the *G9a* cKO, although to a lower, often non-significant magnitude. Furthermore, there appears to be a potentially unique role of GLP in oocyte DNA methylation, although this is likely to be minor compared to its function compensatory to loss of G9A.

To assess this possibility in more detail, we analysed the regions of the genome affected by DNA methylation changes. While hypermethylated DMRs show similar overlap with genic and intergenic regions, hypomethylated DMRs are enriched in genic regions (**Supplemental Fig. S4E**). This is expected, as hypomethylated DMRs are found in regions that are methylated in control oocytes and it is well established that DNA methylation localises predominantly to transcribed genes in oocytes (Veselovska et al. 2015). To assess how DNA methylation changes correlate with underlying histone modifications, we tested the overlap of DMRs with H3K9me2, H3K27me3 and H3K4me3 (**Fig. 4F**). The majority of DMRs did not overlap regions with H3K9me2 and there was no clear distinction between hyper- and hypomethylated DMRs, indicating that most DNA methylation changes in cKO oocytes are unlikely to be a direct consequence of loss of H3K9me2. Interestingly, hypermethylated DMRs were strongly enriched for H3K27me3, compared to hypomethylated DMRs. Importantly, this enrichment was not seen for H3K4me3, although both are enriched in regions of the genome lacking DNA methylation. Because H3K27me3 and DNA methylation are mutually exclusive, this result may indicate a localised redistribution of the two marks in cKOs.

As transcription is linked to the deposition of gene-body DNA methylation in the mouse oocyte, we analysed the correlation between expression and DNA methylation changes. Consistently, expression changes of DEGs positively correlated with gene-body DNA methylation changes in *G9a-Glp* cDKO oocytes (**Fig. 4G**). In contrast, when evaluating genes with differential methylation, the correlation was much weaker (**Supplemental Fig. S5F**). These trends were similar in *G9a* cKO oocytes (**Supplemental Fig. S5G,H**). This shows that the transcriptional changes observed in *G9a-Glp* cDKO oocytes impact the associated genic DNA methylation; however, this association does not explain most of the DNA methylation changes in *G9a-Glp* cDKO oocytes, thus other mechanisms likely underlie the majority of changes observed.

Since G9A and GLP can interact with UHRF1 (Kim et al. 2009; Ferry et al. 2017), we also investigated the overlap of DMRs with regions that are hypomethylated in *Uhrf1* cKO oocytes (Maenohara et al. 2017). We compared the overlap of *G9a* cKO and *G9a-Glp* cDKO hypomethylated DMRs with regions hypomethylated in *Uhrf1* cKO oocytes, random probes and random probes that are highly methylated in control oocytes, as these are the regions most likely affected by loss of methylation. We observed that both *G9a* cKO and *G9a-Glp* cDKO hypomethylated DMRs strongly overlap with *Uhrf1* hypomethylated regions compared to random and random-methylated probes (**Fig. 4H**). These data suggest that loss of G9A and GLP may disturb a subset of UHRF1-mediated methylation. In summary, our analysis suggests that genic DNA methylation gains seen in *G9a* cKO and *G9a-Glp* cDKO oocytes occur largely as a consequence to gene de-repression, while losses of DNA methylation may be linked to impaired UHRF1-mediated *de novo* DNA methylation activity. These findings highlight that G9A and GLP are integral to several parallel molecular processes.

## Discussion

Our study shows that GLP can partially compensate for loss of G9A and additionally has a G9A- independent role in oogenesis (summarized in **Fig. 5**). We find that GLP is required for oocyte maturation and developmental competence. Few embryos derived from *Glp* cKO oocytes implant and those that do die mid-gestation. In contrast, embryos from *G9a* cKO oocytes are less affected and some develop normally into healthy pups. This difference in severity is also reflected at a molecular level in oocytes, with a marked de-repression of gene expression which is correlated with increased protein abundance and gains in genic DNA methylation. Notably, the differences between the genotypes are apparently independent of GLP’s function as an H3K9me2 methyltransferase, as H3K9me2 was equally ablated in all cKOs as measured by IF and ChIP-seq. However, the loss of H3K9me1 only in the *Glp* cKO and *G9a-Glp* cDKO oocytes suggests that GLP’s effect as a transcriptional repressor could be mediated in part through H3K9me1. Importantly, transcriptional changes do not explain all observed changes in DNA methylation with a marked overlap of regions that lose DNA methylation with those that are dependent on UHRF1, suggesting G9A and GLP may play a role in directing some UHRF1-facilitated DNA methylation.

**Figure 5.**
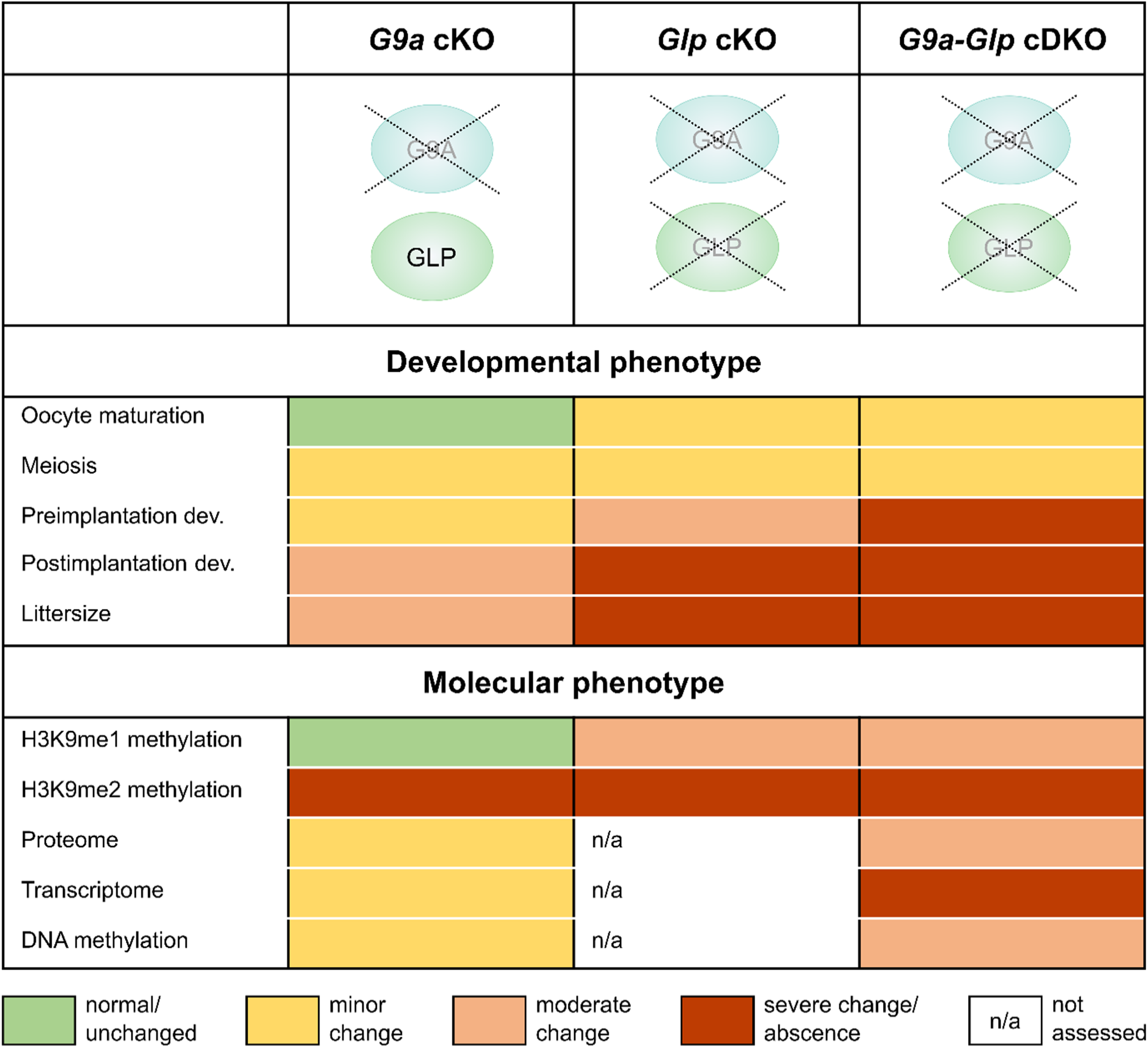
Summary of the developmental and molecular phenotypes of *G9a* cKO, *Glp* cKO and *G9a- Glp* cDKO oocytes demonstrating the distinct roles of GLP and G9A in the oocyte.

To consider the molecular mechanisms underlying the observed phenotypes, it is important to appreciate that G9A and GLP are proteins with multiple functions. Although best known as histone methyltransferases, they can methylate and alter the function of non-histone proteins (Shinkai and Tachibana 2011; Scheer and Zaph 2017). Nevertheless, the strong correlation we observed between changes in transcript and protein abundance in *G9a* and *G9a/Glp* cKO oocytes argues that the changes in protein abundance are likely attributable to altered genomic regulation rather than G9A/GLP directly modulating protein stability.

Loss of G9A and GLP resulted in transcriptional upregulation, supporting previous reports that G9A/GLP act as repressors (Tachibana et al. 2002; Tachibana et al. 2005). It is unlikely that this can be attributed to loss of H3K9me2, because of the small number of genes that were upregulated despite almost complete ablation of H3K9me2, and because of the significant transcriptional differences between the *G9a* cKO and *G9a-Glp* cDKO despite similar deficits in H3K9me2. Alternatively, the effects may be mediated indirectly by G9A/GLP modulating activity of transcriptional repressors or through other chromatin changes, such as H3K9me1 or potentially H3K27me2. Further elucidation of whether changes in these repressive marks may be linked to transcriptional de-repression remains a challenge, due to a lack of robust antibodies for ULI-ChIP-seq. We also observed upregulation of some developmental transcription factors, suggesting that some gene expression changes reflect illegitimate expression of such factors in oocytes. The extensive transcriptional de-repression in the *G9a-Glp* cDKO is associated with impaired developmental capacity of the oocyte, resulting in decreased oocyte maturation, impaired fertilization and abnormal embryo development.

G9A and GLP have been linked to DNA methylation in multiple studies. In the embryo, loss of G9A leads to hypomethylation of a subset of CGI promoters (Auclair et al. 2016). In the oocyte, we and others do not see widespread hypomethylation (Au Yeung et al. 2019) but, instead, local sites of hypo- and hypermethylation. A variety of mechanisms could underpin these changes. In the oocyte, *de novo* DNA methylation requires transcription (Kobayashi et al. 2012; Veselovska et al. 2015). The transcriptional changes in *G9a* cKO and *G9a-Glp* cDKO oocytes correlate with DNA methylation changes, indicating that upregulated gene expression is responsible for some of the hypermethylation observed. The significant localisation of *G9a* cKO and *G9a-Glp* cDKO de-repressed genes and hypermethylated regions with H3K27me3 suggest that the loss of G9A/GLP may impair the repressive chromatin landscape in a subset of regions the oocyte, which could reflect its action in depositing H3K9me1/2, modulating levels of H3K27me2, or activity as a transcriptional repressor. These mechanisms warrant further study, but possibly in other cell contexts because determining cause versus effect would be challenging in the oocyte.

A large subset of hypomethylated DMRs overlapped with regions that are hypomethylated in *Uhrf1* cKO oocytes (Maenohara et al. 2017). UHRF1 function has been associated with G9A and GLP in several ways, either by direct interaction (Kim et al. 2009) or indirect recruitment through H3K9me2/3 or LIG1 (Rothbart et al. 2012; Rothbart et al. 2013; Ferry et al. 2017). However, it is not clear how these mechanisms apply in the oocyte because of the absence of DNA replication. Although UHRF1 appears to be required for some genomic DNA methylation, loss of DNMT1 has only minor effects, largely associated with its role in ensuring symmetric methylation of *de novo* methylated CpGs (Shirane et al. 2013; Maenohara et al. 2017). There are likely to be other mechanisms besides transcriptional regulation and UHRF1 interactions relevant in the *G9a* cKO and *G9a-Glp* cDKO oocytes. These include the possibility of a direct interaction of G9A and GLP with DNMT proteins; for example, G9A and GLP can dimethylate DNMT3A at lysine 44, and methylated DNMT3A can be bound by MPP8, which in turn binds automethylated GLP, resulting in a DNMT3A-MPP8-GLP silencing complex (Chang et al. 2011).

In line with previous studies (Au Yeung et al. 2019), we find that G9A is essential for H3K9me2 establishment in the oocyte, but not H3K9me1. While H3K9me2 depends on the presence of G9A, the persistence of GLP in the *G9a* cKO was sufficient to establish H3K9me1. Notably, loss of G9A and GLP did not result in a complete ablation of H3K9me1, indicating that other methyltransferases, such as PRDM3 and PRDM16, may be active in the oocyte. It remains unclear whether the decrease in H3K9me1 may be at least partially causative for the transcriptional and DNA methylation changes. Thus far, H3K9me1 has not been associated with transcriptional repression or *de novo* DNA methylation, but it is under-characterised relative to H3K9me2/3 due to a lack of validated, high quality ChIP-grade antibodies.

Our results indicate that GLP can compensate for loss of G9A in the oocyte, but also has unique roles. G9A and GLP are thought to function as heterodimers *in vivo*. Because G9A is unstable on its own, it is technically challenging to create a model with intact G9A in the absence of GLP. When comparing transcription, protein and DNA methylation data from *G9a* cKO and *G9a-Glp* cDKO oocytes, some genes, proteins and genomic regions are progressively affected while others only in *G9a-Glp* cDKO oocytes. The possibility of a unique, G9A-independent role for GLP is supported by a recent study which reported that GLP interacts with PRC2 in zygotes to establish H3K27me2 in the paternal pronucleus (Meng et al. 2020). In our models, the unique function was especially apparent in the transcriptome, where genes with the greatest fold-change where unique to the *G9a-Glp* cDKO. The upregulation seen in most DEGs was reflected in upregulation of protein abundance in *G9a-Glp* cDKO oocytes, indicating that although only a relatively small proportion of genes is derepressed upon loss of G9A and GLP, the transcriptional changes are likely to have a functional impact in the oocyte. Indeed, we saw changes in gene expression and protein abundance of several genes related to oocyte maturation and fertilization. But the elevated or inappropriate expression of genes and corresponding proteins may be more deleterious than down-regulation, as very few genes are likely to be haploinsufficient in oocytes.

Taken together, our results highlight GLP as a multi-functional repressive protein required for the appropriate establishment of the oocyte transcriptome, epigenome and proteome. Consequently, GLP is critical for the developmental capacity of the oocyte, independent of G9A.

## Materials and methods

### Sample collections

All mice used in this study were bred and maintained in the Babraham Institute Biological Support Unit. Ambient temperature was ∼19-21°C and relative humidity 52%. Lighting was provided on a 12 hour light: 12 hour dark cycle including 15 min ‘dawn’ and ‘dusk’ periods of subdued lighting. After weaning, mice were transferred to individually ventilated cages with 1-5 mice per cage. Mice were fed CRM (P) VP diet (Special Diet Services) *ad libitum* and received seeds (e.g. sunflower, millet) at the time of cage-cleaning as part of their environmental enrichment. All experimental procedures were performed under licences issued by the Home Office (UK) in accordance with the Animals (Scientific Procedures) Act 1986 and were approved by the Animal Welfare and Ethical Review Body at the Babraham Institute.

Samples were collected from cKO mice carrying a *Zp3*-Cre driver in addition with floxed alleles for *G9a* (Sampath et al. 2007), *Glp* (Schaefer et al. 2009) or both. Oocytes and embryos collected in M2 medium (Sigma-Aldrich, M7167) unless stated otherwise. GV and MII oocytes for IF analysis were collected from adult mice aged ∼12 weeks and fixed in 2% PFA for 15 minutes. MII oocytes were collected after superovulation. Oocytes used for ChIP-seq, BS-seq and RNA-seq were collected from ovaries of 22-26 day-old mice using a collagenase/trypsin digest. For ChIP-seq, 300 GV oocytes were collected in nuclear lysis buffer and pooled from two to four mice for each replicate. For BS-seq and RNA-seq, replicates were stored in RLT+ buffer (Qiagen). Each replicate comprised all oocytes collected from one mouse (75-200 GV oocytes). For whole-proteome analysis oocytes were collected from ovaries of 22-26 day-old mice. To avoid contamination with proteins/peptides from the medium, oocytes were collected by manual dissection of ovaries in protein-free L15 medium (Thermo Fisher Scientific, 31415029). ∼200 GV oocytes were collected from two mice in parallel, washed 3 times in PBS with 1x cOmplete Protease Inhibitor Cocktail (Merck, 11697498001).

Preimplantation embryos were collected after natural mating of control and cKO females with C57BL/6Babr WT males at E0.5 or E3.5. Implantation was determined by counting the number of implantation sites in uteri at E6.5 after timed mating. To score postimplantation development, embryos were collected after natural mating at E8.5 and E12.5. For embryo culture, female mice were superovulated and embryos dissected in M2 medium at E1.5 after natural mating with WT males. Fertilized embryos (1-cell and 2-cell stage) were selected for culture in M16 medium (Sigma-Aldrich, MR-016-D) under mineral oil (Sigma-Aldrich, M8410) at 37°C and 5% CO_2_ for 3 days, and developmental progress recorded each day.

### Analysis of maturation stage

GV oocytes from 12 week-old females were stained with DAPI and scored according to their maturation stage. Absence of a ring around the nucleolus was counted as “NSN”, a partial ring as “intermediate” and a full ring “SN”. Between 116 and 247 oocytes collected from several different females were analysed per genotype (Number of mice: Control=5, G9a cKO=3, Glp cKO=2, G9a-Glp cDKO=3).

### Immunofluorescence analysis

IF was performed after antibody staining as previously described (Santos et al. 2003). Primary antibodies are listed in **Supplemental Table S5**. MII oocytes were stained with antibodies against α- tubulin (spindle), pan-histone (chromatin) and DAPI (heterochromatic DNA). Between 46 and 57 MII oocytes from 4 to 7 mice were analysed per genotype. For H3K9me1, H3K9me2, H3K9me3, 5-mC and 5-hmC two or three replicate experiments were performed, each with 10-15 oocytes from different litters, to control for batch effects. Samples were analysed on a Zeiss LSM780 confocal microscope (63x oil-immersion objective). Spindle conformation and chromosome alignment of MII oocytes was scored using categories illustrated in **Supplemental Fig. S1C**. For analysis of H3K9me and DNA methylation Z-stacks of single optical sections were captured and semi-quantification of fluorescence intensity was performed using Volocity 6.3 (Improvision).

### LC-MS proteome analysis

Oocytes were lysed in 20µl dissolution buffer containing 100mM triethylammonium bicarbonate (Sigma, T4708) and 0.1% Sodium Dodecyl Sulfate (SDS), followed by water bath sonication and boiling at 90°C for 5min. Proteins were reduced with tris-2-carboxyethyl phosphine (ΤCEP, Sigma) for 1h at 60°C at a final concentration of 5mM, followed by cysteine blocking for 10min at room temperature using methyl methanethiosulfonate (MMTS, Sigma) at final concentration of 10mM. Samples were digested overnight at 37°C with trypsin (Pierce #90058) and the next day peptides were labelled with TMT11plex reagents (0.4mg per sample) according to manufacturer’s instructions (Thermo Scientific). To quench the reaction, 3µl of 5% hydroxylamine (Thermo Scientific) was added for 15min and samples combined and dried with centrifugal vacuum concentrator. The TMT mix was fractionated with Reversed-Phase spin columns at high pH (Pierce #84868). Nine fractions were collected using different elution solutions in the range of 5–50% ACN and were analysed on a Dionex UltiMate 3000 UHPLC system coupled with the nano-ESI Fusion-Lumos (Thermo Scientific) mass spectrometer. Samples were loaded on the Acclaim PepMap 100, 100μm × 2cm C18, 5μm, 100Ȧ trapping column with the ulPickUp injection method at loading flow rate 5μl/min for 10 min. For peptide separation, the EASY-Spray analytical column 75μm × 25cm, C18, 2μm, 100 Ȧ column was used for multi-step gradient elution. Full scans were performed in the Orbitrap in the range of 380-1500 m/z at 120K resolution and peptides isolated in the quadrupole with isolation window 1.2Th, HCD collision energy 38% and resolution 50K. Raw data were processed with the SequestHT search engine in Proteome Discoverer 2.1 software and searched against a Uniprot database containing mouse reviewed entries. The parameters for the SequestHT node were: Precursor Mass Tolerance 20ppm, Fragment Mass Tolerance 0.02Da, Dynamic Modifications were Oxidation of M (+15.995Da), Deamidation of N, Q (+0.984Da) and Static Modifications were TMT6plex at any N-Terminus, K (+229.163Da) and Methylthio at C (+45.988Da). The consensus workflow included TMT signal-to-noise (S/N) calculation and the level of confidence for peptide identifications was estimated using the Percolator node with decoy database search. Strict FDR was set at q-value<0.01. For downstream data analysis, the R package qPLEXanalyzer was used (Papachristou et al. 2018). The mass spectrometry proteomics data have been deposited to the ProteomeXchange Consortium via the PRIDE partner repository (Perez-Riverol et al. 2019) with the dataset identifier PXD030265.

### Preparation of sequencing libraries

ChIP-seq libraries were prepared using ULI-nChIP as previously described (Hanna et al. 2018b). Antibodies were added at 250ng/reaction for both anti-H3K9me2 (mouse monoclonal, Abcam, ab1220) and anti-IgG (rabbit polyclonal, Diagenode, EB-070-010). Library preparation was completed with a MicroPlex Library Preparation kit v2 (Diagenode) with Sanger 8-base indices for multiplexing. Relative enrichment over input was quantified using the library concentrations determined by Bioanalyzer High Sensitivity DNA Analysis (Agilent). Low input bisulphite (BS)-seq libraries were prepared by post-bisulphite adapter tagging as previously described (Hanna et al. 2018b). RNA-seq libraries were prepared as described (Hanna et al. 2019).

### Sequencing and data processing

Libraries were sequenced on Illumina MiSeq, HiSeq2500 and NextSeq500 systems. ChIP-seq libraries were sequenced to an average of 57 million paired-end reads of 75 bp read-length (**Supplemental Table S5**). BS-seq libraries were sequenced to an average of 16 million paired-end reads for *G9a* cKO and 26 million reads for *G9a-Glp* cDKO oocytes of 100-125 bp read-length. RNA-seq libraries were sequenced to an average single read number of 1.9 million for *G9a* cKO and *G9a-Glp* cDKO oocytes of 50 bp read-length. Raw fastq files were processed with Trim Galore, then mapped to the mouse GRCm38 genome. Mapping of ChIP-seq data was done using Bowtie2, RNA-seq data by Hisat2 v2.1.0 guided by known splice sites, BS-seq data with Bismark v0.19.0.

### Sequencing data analysis

Sequencing data analysis was conducted using SeqMonk (https://www.bioinformatics.babraham.ac.uk/projects/seqmonk/). For ChIP-seq analysis, 10kb running windows (*N*=272,566) were quantified as reads per kilobase per million (RPKM). Windows were filtered to exclude mapping artefacts, defined as RPKM>6 in at least one replicate set of 10% input libraries (*N*=408). H3K9me2 enrichment was defined as Log_2_RPKM>2.5 in d25 GV oocytes (*N*=34,192). A set of random windows was sampled from all 10kb windows (*N*=35,000). H3K9me2- enriched and random windows were then merged with adjacent windows within 10kb, resulting in 12,154 H3K9me2-enriched domains and 26,077 random domains. Genic and intergenic regions were defined as overlapping or not overlapping oocyte transcripts, respectively, and promoters were defined as +/-500bp around transcription start sites of oocyte transcripts (Veselovska et al. 2015). CGIs were defined as previously described (Illingworth et al. 2008). Oocyte transcription levels were categorized into not expressed (FPKM<0.1), low expressed (FPKM 0.1-1) and high expressed (FPKM>1) using published data (Veselovska et al. 2015).

BS-seq data were analysed using a tile-based approach of 100 CpGs for each consecutive genome window, ensuring equal CpG content in all windows analysed. Methylation values were quantified using the bisulphite-sequencing pipeline quantification, which calculates per-base methylation percentages and averages these within each window. Filters were applied to ensure a minimum coverage of ≥10 observed cytosines per window. Only windows with this minimal coverage in all samples were taken into account, allowing assessment of 86.2% of 100 CpG windows (*N*=188,433). Differentially methylated regions (DMRs) were defined by statistical comparison of DNA methylation levels for each 100 CpG window between control and *G9a* cKO or control and *G9a-Glp* cDKO oocytes using the EdgeR function in SeqMonk. To assess overlap of DMRs with genomic features, CGI and oocyte gene annotations were used from (Veselovska et al. 2015). DMRs were merged to form differentially methylated domains (DMDs). Hierarchical clustering analysis of DMDs was performed in R using Euclidean distance and Ward’s agglomeration method as implemented by the package pheatmap.

For analysis of RNA-seq data, expression of oocyte genes (Veselovska et al. 2015) was quantified using log transformed read count quantitation per million reads. Differential gene expression was analysed using DESeq2 followed by filtering of genes with Log_2_FC>1.5. Clustering analysis of DEGs was performed in R using Euclidean distance and Ward’s agglomeration method. Relative enrichment of differentially abundant proteins amongst DEGs was displayed using the barcode method from the Limma R package. The ranked list of expression changes only considered genes corresponding to proteins identified in the proteome. Spearman’s rank correlation coefficient between DNA methylation difference and expression log_2_FC was used to interrogate the relationship between DNA methylation and transcriptional changes.

Sequencing data were submitted to the GEO under accession number GSE191026. Publicly available ChIP-seq and BS-seq data under accessions GSE93941, DRA005849 were used to develop our studies.

### Statistical analysis

Statistical analysis was conducted in Graphpad Prism, R and VassarStats. SN proportion was compared using two-way ANOVA (Stage x Genotype) with Šídák’s multiple comparisons test. Proportion of embryo stage at E3.5, number of implantation sites at E6.5 and mean fluorescence intensity of confocal IF images were analysed using one-way ANOVA with Tukey’s posthoc test. Litter sizes were compared using a nested one-way ANOVA with mouse as nested factor. A χ-square test was used to analyse meiosis abnormalities and the frequency of overlap of genic/intergenic regions with DMRs, histone modifications with DMRs, Uhrf1 hypomethylated regions with DMRs and DEGs with histone modifications. *P*-value was adjusted using Bonferroni correction to control for multiple comparisons.

## Acknowledgements

We would like to Kristina Tabbada and Nicole Forrester of the Sequencing, Simon Walker of the Imaging, Simon Andrews and Felix Krueger of the Bioinformatics facilities, and members of the Biological Support Unit at the Babraham Institute for excellent support. Work in G.K.’s lab was funded by the UK Biotechnology and Biological Sciences Research Council (BBS/E/B/000C0423) and Medical Research Council (MR/K011332/1, MR/S000437/1); H.D. was supported by a Fondecyt Postdoctorado fellowship (3200676); C.H. was supported by a Next Generation Fellowship from the Centre for Trophoblast Research. Part of the work in C.D’S.’s lab was funded by CRUK core grant (A29580) awarded to the Cambridge Institute.

## Author contributions

C.H., H.D. and G.K. conceptualised the study. H.D., F.S., E.K.P. and C.H. collected the data. H.D., C.H., J.C-F., and K.K. performed data analysis. H.D., C.H. and J.C.-F. generated manuscript figures. H.D. drafted the manuscript, with input from C.H., J.C.-F., E.K.P., C.D’S. and G.K.

## Supplementary Figures

**Figure S1:**
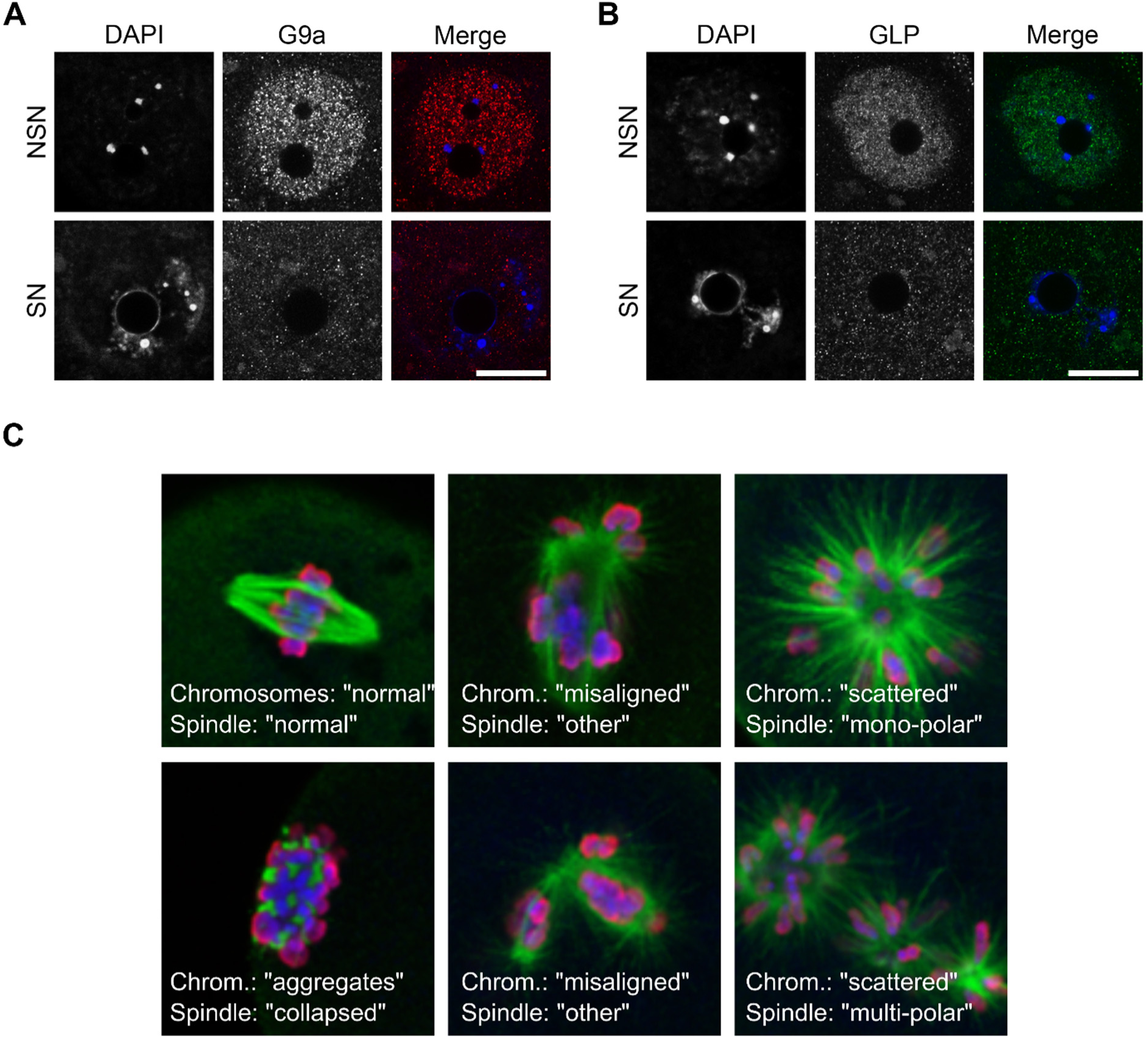
G9A and GLP expression in the oocyte and their effects on meiosis. **A, B)** Representative images showing DNA stained with DAPI, and G9A expression **(A)** and GLP expression **(B)** revealed by IF in NSN and SN control oocytes. The scale bar represents 20µm. **C)** Example images showing meiotic abnormalities observed in MII oocytes. The spindle is stained with an anti-α-tubulin antibody (green) and the chromatin with DAPI (blue) and anti-pan-histone (red).

**Figure S2:**
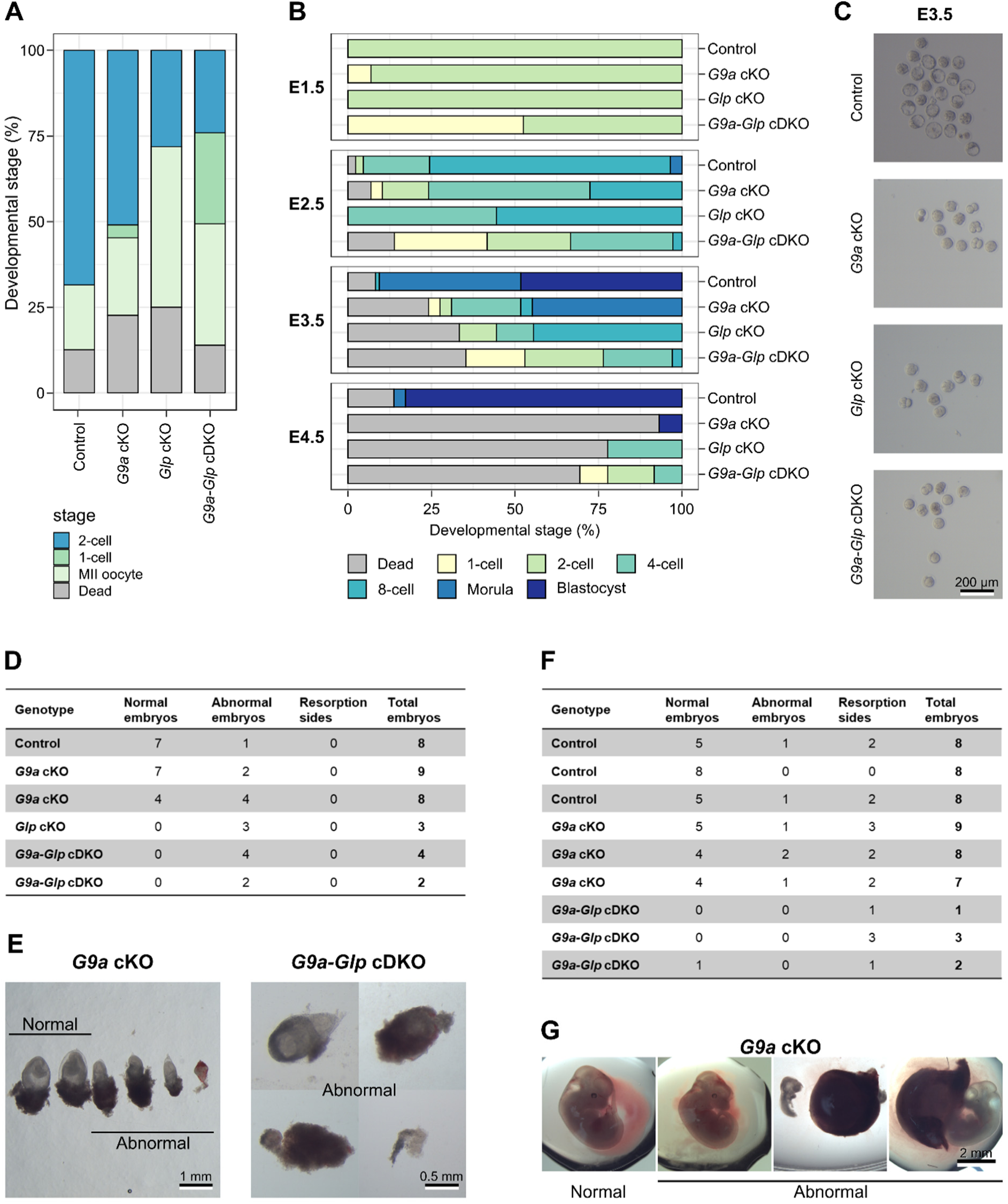
I*n vitro* preimplantation and *in vivo* postimplantation development of embryos lacking maternal G9A and/or GLP. **A)** Stacked barchart showing developmental stage of embryos collected at E1.5 after superovulation and natural mating with C57Bl6/Babr WT males. . One-way ANOVA with Tukey’s posthoc test: 2-cell stage Control vs G9a-Glp cDKO *P = 0.0068\*\**. **B)** Stacked barcharts showing developmental stage of embryos cultured *in vitro* after natural mating on E1.5, E2.5, E3.5 and E4.5. Number of litters/total number embryos: Control = 3/86, *G9a* cKO = 3/29, *Glp* cKO = 1/9, *G9a-Glp* cDKO = 4/40. One-way ANOVA with Tukey’s posthoc test: morula stage Control vs *G9a* cKO *P = 0.0006\*\*\**. **C)** Images showing examples of embryos cultured in vitro on E3.5. **D, E)** Table showing embryo development at E8.5 **(D)** and E12.5 **(E)** after natural mating of conditional KO females with WT males. Each line represents one female mouse. Indicated are the number of normal embryos, abnormal embryos and resorption sides for each female.

**Figure S3:**
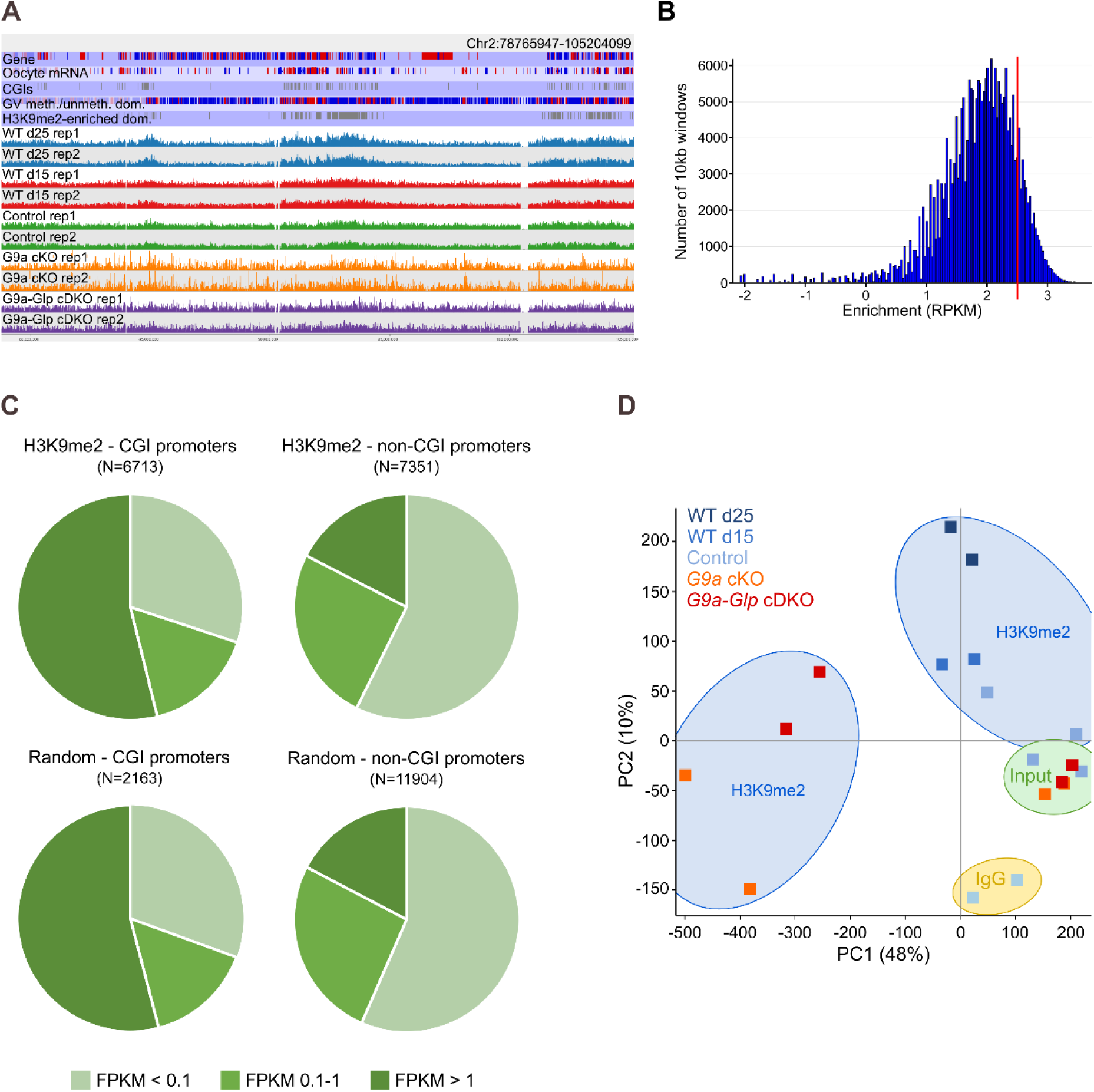
ChIP-seq analysis showing H3K9me2 distribution in control, *G9a* cKO and *G9a-Glp* cDKO oocytes. **A)** Screenshot showing H3K9me2 enrichment and reproducibility of replicates. **B)** Histogram showing number of 10 kb windows with a certain enrichment level in d25 WT oocytes. The red line indicates the threshold set to define H3K9me2 enrichment (RPKM > 2.5). **C)** Pie charts showing overlap of H3K9me2 enriched and random probes with untranscribed (FPKM < 0.1), lowly transcribed (FPKM 0.1- 1) and highly transcribed genes (FPKM > 1). **D)** PCA plot of H3K9me2 (blue shading), input (green shading) and IgG (yellow shading) ChIP-seq libraries, comparing WT (d15 and d25), control, *G9a* cKO and *G9a-Glp* cDKO samples.

**Figure S4:**
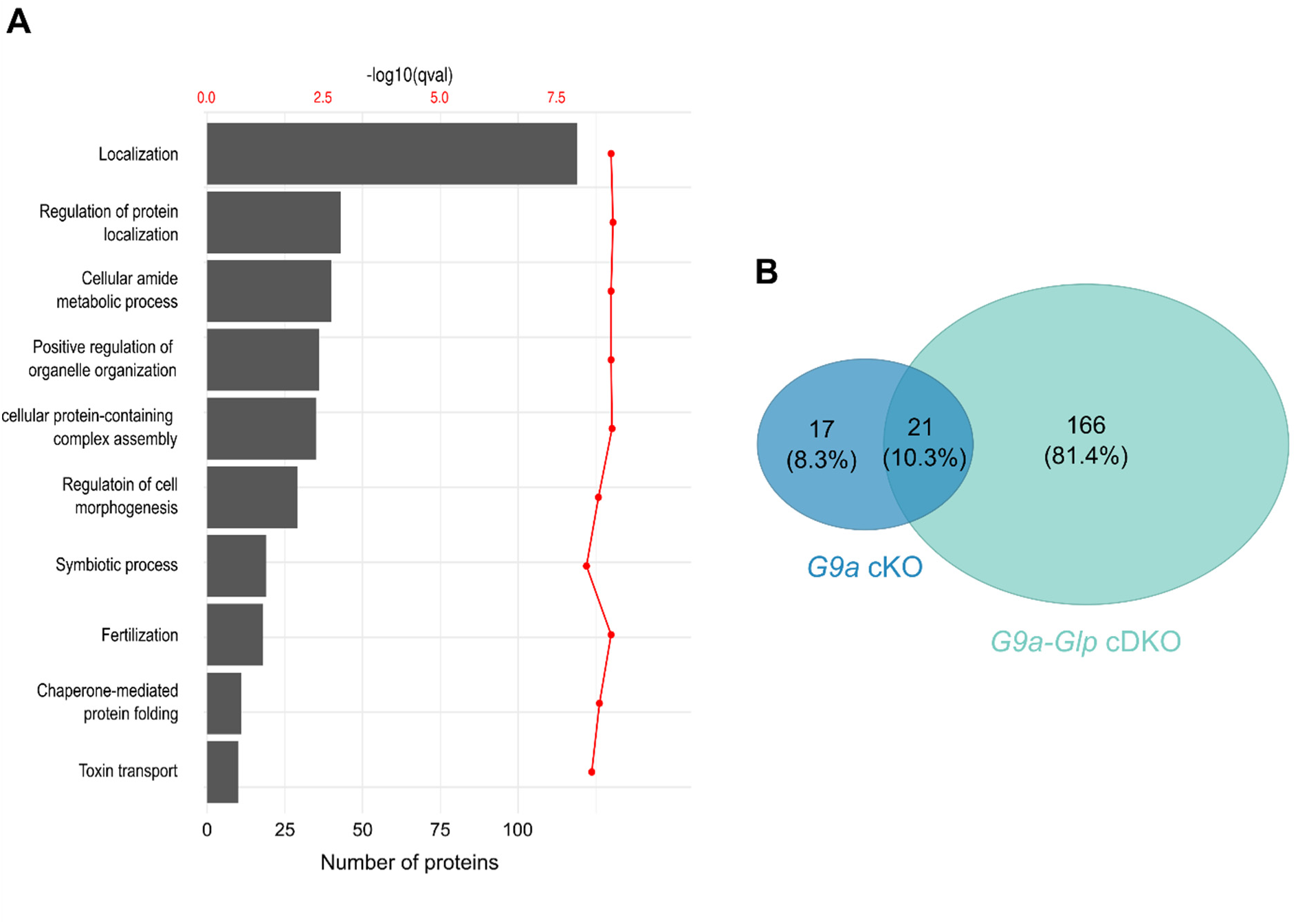
Proteome and transcriptome analysis of *G9a* cKO and *G9a-Glp* cDKO oocytes with corresponding littermate controls. **A)** GO analysis of the oocyte proteome. **B)** Venn diagram of significant changing proteins in *G9a* cKO and *G9a-Glp* cDKO oocytes.

**Figure S5:**
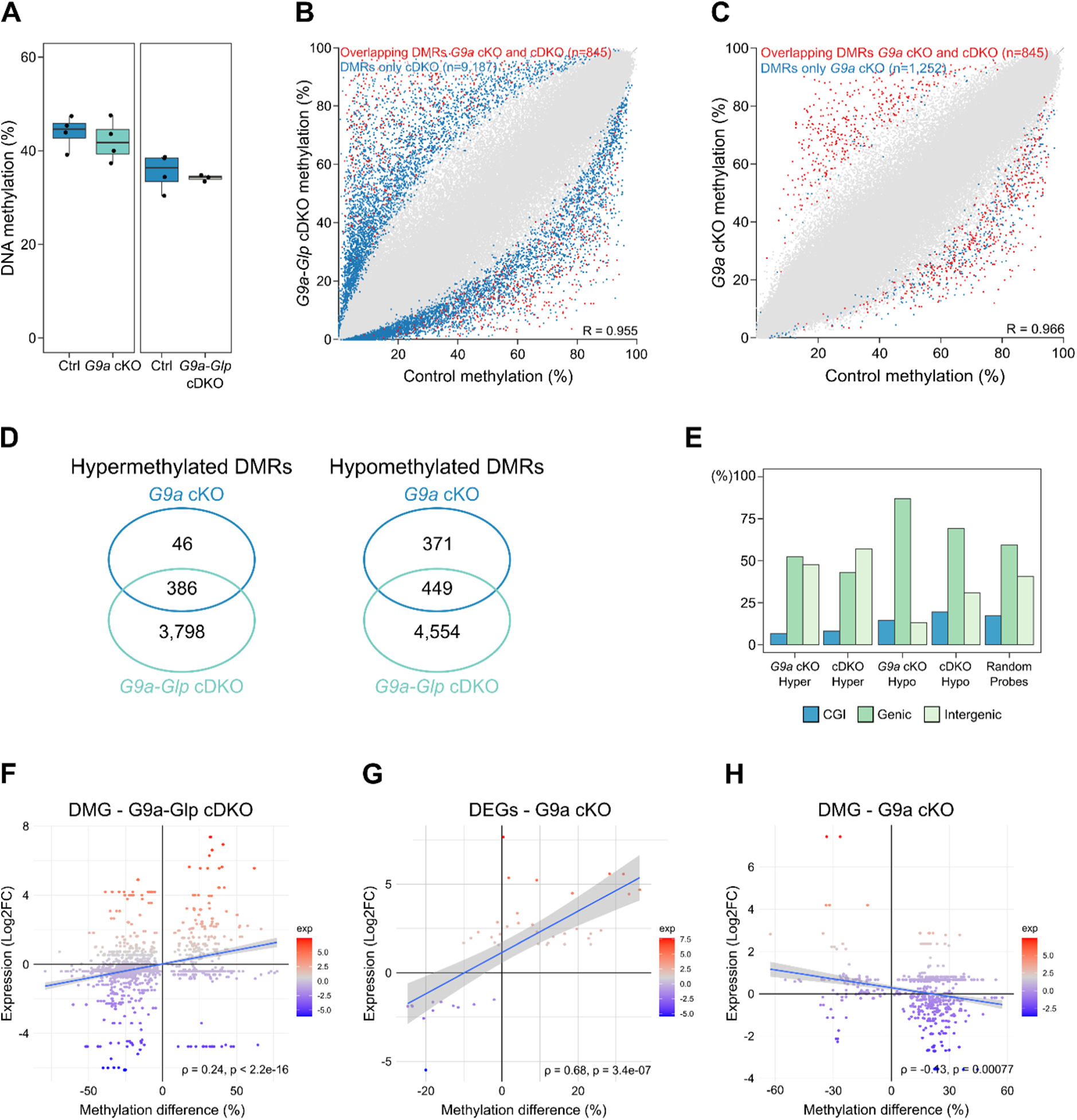
DNA methylation analysis of *G9a* cKO and *G9a-Glp* cDKO oocytes with corresponding littermate controls. **A)** Box-whisker plots showing global DNA methylation levels using average methylation levels of 100 CpG windows. Ctrl *vs G9a* cKO *P* = 0.5378; Ctrl vs *G9a-Glp* cDKO *P* = 0.6077 **B,C)** Scatterplot showing methylation levels of control vs *G9a-Glp* cDKO **(B)** and control vs *G9a* cKO oocytes **(C)**. Each dot represents the methylation values of a 100 CpG window. Unique DMRs are highlighted in blue and DMRs shared between the two genotypes in red. **D)** Venn diagrams, showing overlap of hypermethylated and hypomethylated DMRs between *G9a* cKO and *G9a-Glp* cDKO oocytes. **E)** Barchart showing percentage of DMRs and random probes overlapping CGIs, genic and intergenic regions. χ-square comparing genic and intergenic DMRs: G9a cKO *adj. P = 0.026\**; all others *P < 0.0001\*\*\**. **F-H)** Scatterplots showing link between methylation and expression changes of differentially methylated genes (DMG) in *G9a-Glp* cDKO oocytes **(F)**, differentially expressed genes (DEGs) in *G9a* cKO oocytes **(G)** and DMGs in *G9a* cKO oocytes **(H)**. Expression levels are indicated by the colour scale. Spearman correlation indicated in plot.

**Figure S6:**
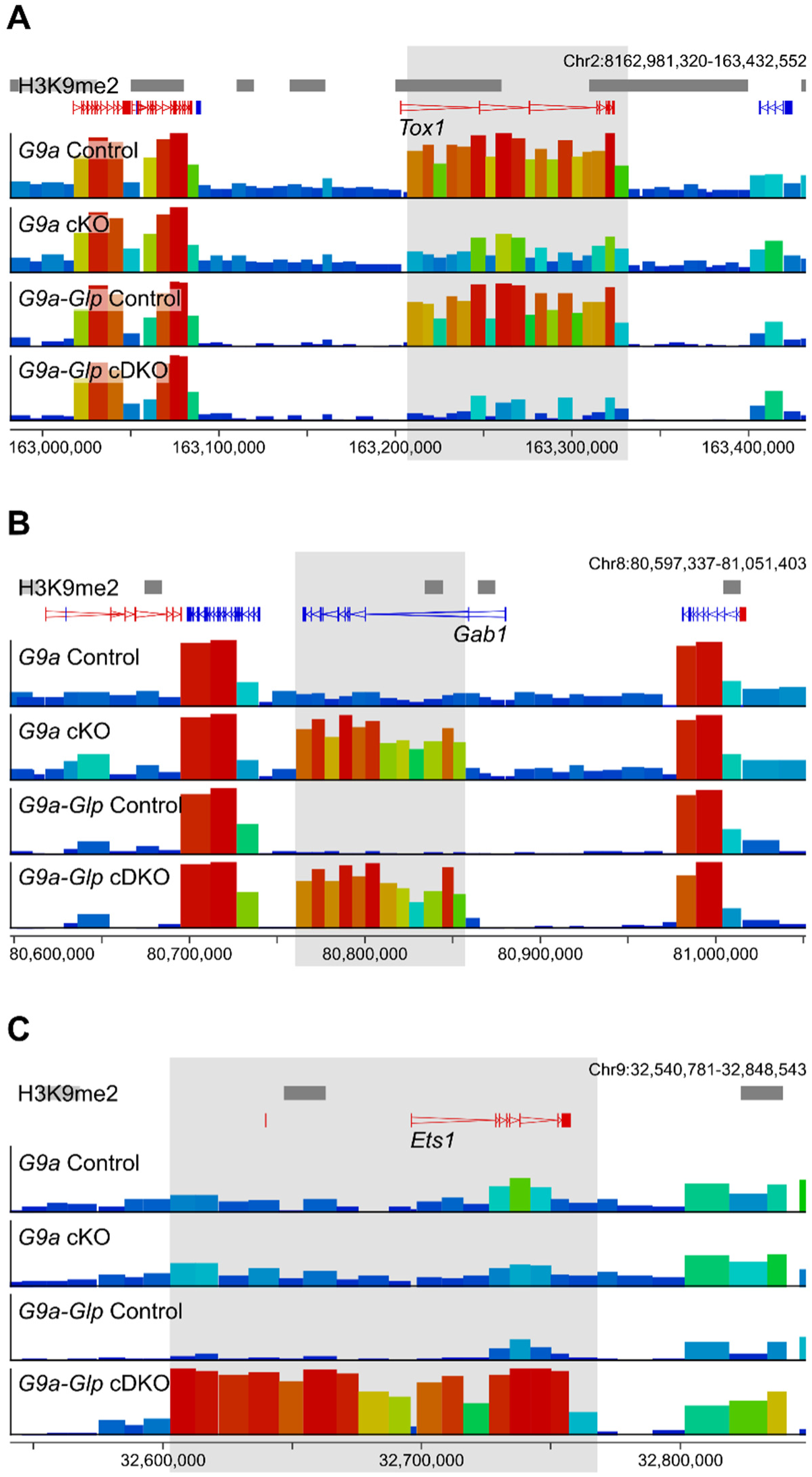
Genome screenshots showing examples of differentially methylated domains. Shown are: a region that is hypomethylated in both *G9a* cKO and *G9a-Glp* cDKO oocytes **(A)**, a region hypermethylated in both *G9a* cKO and *G9a-Glp* cDKO oocytes **(B),** and a region uniquely hypermethylated in *G9a-Glp* cDKO oocytes **(C)**.

## References

Andreu-Vieyra CV, Chen R, Agno JE, Glaser S, Anastassiadis K, Stewart AF, Matzuk MM. 2010. MLL2 is required in oocytes for bulk histone 3 lysine 4 trimethylation and transcriptional silencing. PLoS Biol 8.

Au Yeung WK, Brind’Amour J, Hatano Y, Yamagata K, Feil R, Lorincz MC, Tachibana M, Shinkai Y, Sasaki H. 2019. Histone H3K9 Methyltransferase G9a in Oocytes Is Essential for Preimplantation Development but Dispensable for CG Methylation Protection. Cell reports 27: 282–293 e284.

Auclair G, Borgel J, Sanz LA, Vallet J, Guibert S, Dumas M, Cavelier P, Girardot M, Forne T, Feil R et al. 2016. EHMT2 directs DNA methylation for efficient gene silencing in mouse embryos. Genome Res 26: 192–202.

Bittencourt D, Lee BH, Gao L, Gerke DS, Stallcup MR. 2014. Role of distinct surfaces of the G9a ankyrin repeat domain in histone and DNA methylation during embryonic stem cell self-renewal and differentiation. Epigenetics Chromatin 7: 27.

Chang Y, Sun L, Kokura K, Horton JR, Fukuda M, Espejo A, Izumi V, Koomen JM, Bedford MT, Zhang X et al. 2011. MPP8 mediates the interactions between DNA methyltransferase Dnmt3a and H3K9 methyltransferase GLP/G9a. Nat Commun 2: 533.

Collins RE, Northrop JP, Horton JR, Lee DY, Zhang X, Stallcup MR, Cheng X. 2008. The ankyrin repeats of G9a and GLP histone methyltransferases are mono- and dimethyllysine binding modules. Nat Struct Mol Biol 15: 245–250.

Dong KB, Maksakova IA, Mohn F, Leung D, Appanah R, Lee S, Yang HW, Lam LL, Mager DL, Schubeler D et al. 2008. DNA methylation in ES cells requires the lysine methyltransferase G9a but not its catalytic activity. EMBO J 27: 2691–2701.

Eymery A, Liu Z, Ozonov EA, Stadler MB, Peters AH. 2016. The methyltransferase Setdb1 is essential for meiosis and mitosis in mouse oocytes and early embryos. Development doi:10.1242/dev.132746.

Ferry L, Fournier A, Tsusaka T, Adelmant G, Shimazu T, Matano S, Kirsh O, Amouroux R, Dohmae N, Suzuki T et al. 2017. Methylation of DNA Ligase 1 by G9a/GLP Recruits UHRF1 to Replicating DNA and Regulates DNA Methylation. Mol Cell 67: 550–565 e555.

Guibert S, Forne T, Weber M. 2012. Global profiling of DNA methylation erasure in mouse primordial germ cells. Genome Res 22: 633–641.

Hanna CW, Demond H, Kelsey G. 2018a. Epigenetic regulation in development: is the mouse a good model for the human? Hum Reprod Update 24: 556–576.

Hanna CW, Perez-Palacios R, Gahurova L, Schubert M, Krueger F, Biggins L, Andrews S, Colome-Tatche M, Bourc’his D, Dean W, et al. 2019. Endogenous retroviral insertions drive non-canonical imprinting in extra-embryonic tissues. Genome biology 20: 225.

Hanna CW, Taudt A, Huang J, Gahurova L, Kranz A, Andrews S, Dean W, Stewart AF, Colome-Tatche M, Kelsey G. 2018b. MLL2 conveys transcription-independent H3K4 trimethylation in oocytes. Nat Struct Mol Biol 25: 73–82.

Illingworth R, Kerr A, Desousa D, Jorgensen H, Ellis P, Stalker J, Jackson D, Clee C, Plumb R, Rogers J et al. 2008. A novel CpG island set identifies tissue-specific methylation at developmental gene loci. PLoS Biol 6: e22.

Jiang Q, Ang JYJ, Lee AY, Cao Q, Li KY, Yip KY, Leung DCY. 2020. G9a Plays Distinct Roles in Maintaining DNA Methylation, Retrotransposon Silencing, and Chromatin Looping. Cell reports 33: 108315.

Kaneda M, Okano M, Hata K, Sado T, Tsujimoto N, Li E, Sasaki H. 2004. Essential role for de novo DNA methyltransferase Dnmt3a in paternal and maternal imprinting. Nature 429: 900–903.

Kim J, Zhao H, Dan J, Kim S, Hardikar S, Hollowell D, Lin K, Lu Y, Takata Y, Shen J et al. 2016. Maternal Setdb1 Is Required for Meiotic Progression and Preimplantation Development in Mouse. PLoS Genet 12: e1005970.

Kim JK, Esteve PO, Jacobsen SE, Pradhan S. 2009. UHRF1 binds G9a and participates in p21 transcriptional regulation in mammalian cells. Nucleic Acids Res 37: 493–505.

Kobayashi H, Sakurai T, Imai M, Takahashi N, Fukuda A, Yayoi O, Sato S, Nakabayashi K, Hata K, Sotomaru Y et al. 2012. Contribution of intragenic DNA methylation in mouse gametic DNA methylomes to establish oocyte-specific heritable marks. PLoS Genet 8: e1002440.

Lan ZJ, Xu X, Cooney AJ. 2004. Differential oocyte-specific expression of Cre recombinase activity in GDF-9-iCre, Zp3cre, and Msx2Cre transgenic mice. Biol Reprod 71: 1469–1474.

Maenohara S, Unoki M, Toh H, Ohishi H, Sharif J, Koseki H, Sasaki H. 2017. Role of UHRF1 in de novo DNA methylation in oocytes and maintenance methylation in preimplantation embryos. PLoS Genet 13: e1007042.

Meng TG, Zhou Q, Ma XS, Liu XY, Meng QR, Huang XJ, Liu HL, Lei WL, Zhao ZH, Ouyang YC et al. 2020. PRC2 and EHMT1 regulate H3K27me2 and H3K27me3 establishment across the zygote genome. Nat Commun 11: 6354.

Papachristou EK, Kishore K, Holding AN, Harvey K, Roumeliotis TI, Chilamakuri CSR, Omarjee S, Chia KM, Swarbrick A, Lim E et al. 2018. A quantitative mass spectrometry-based approach to monitor the dynamics of endogenous chromatin-associated protein complexes. Nat Commun 9: 2311.

Perez-Riverol Y, Csordas A, Bai J, Bernal-Llinares M, Hewapathirana S, Kundu DJ, Inuganti A, Griss J, Mayer G, Eisenacher M, et al. 2019. The PRIDE database and related tools and resources in 2019: improving support for quantification data. Nucleic Acids Res 47: D442–D450.

Peters AH, Kubicek S, Mechtler K, O’Sullivan RJ, Derijck AA, Perez-Burgos L, Kohlmaier A, Opravil S, Tachibana M, Shinkai Y, et al. 2003. Partitioning and plasticity of repressive histone methylation states in mammalian chromatin. Mol Cell 12: 1577–1589.

Pinheiro I, Margueron R, Shukeir N, Eisold M, Fritzsch C, Richter FM, Mittler G, Genoud C, Goyama S, Kurokawa M et al. 2012. Prdm3 and Prdm16 are H3K9me1 methyltransferases required for mammalian heterochromatin integrity. Cell 150: 948–960.

Rathert P, Dhayalan A, Murakami M, Zhang X, Tamas R, Jurkowska R, Komatsu Y, Shinkai Y, Cheng X, Jeltsch A. 2008. Protein lysine methyltransferase G9a acts on non-histone targets. Nat Chem Biol 4: 344–346.

Rice JC, Briggs SD, Ueberheide B, Barber CM, Shabanowitz J, Hunt DF, Shinkai Y, Allis CD. 2003. Histone methyltransferases direct different degrees of methylation to define distinct chromatin domains. Mol Cell 12: 1591–1598.

Rothbart SB, Dickson BM, Ong MS, Krajewski K, Houliston S, Kireev DB, Arrowsmith CH, Strahl BD. 2013. Multivalent histone engagement by the linked tandem Tudor and PHD domains of UHRF1 is required for the epigenetic inheritance of DNA methylation. Genes Dev 27: 1288–1298.

Rothbart SB, Krajewski K, Nady N, Tempel W, Xue S, Badeaux AI, Barsyte-Lovejoy D, Martinez JY, Bedford MT, Fuchs SM, et al. 2012. Association of UHRF1 with methylated H3K9 directs the maintenance of DNA methylation. Nat Struct Mol Biol 19: 1155–1160.

Sampath SC, Marazzi I, Yap KL, Sampath SC, Krutchinsky AN, Mecklenbrauker I, Viale A, Rudensky E, Zhou MM, Chait BT et al. 2007. Methylation of a histone mimic within the histone methyltransferase G9a regulates protein complex assembly. Mol Cell 27: 596–608.

Santos F, Peat J, Burgess H, Rada C, Reik W, Dean W. 2013. Active demethylation in mouse zygotes involves cytosine deamination and base excision repair. Epigenetics Chromatin 6: 39.

Santos F, Zakhartchenko V, Stojkovic M, Peters A, Jenuwein T, Wolf E, Reik W, Dean W. 2003. Epigenetic marking correlates with developmental potential in cloned bovine preimplantation embryos. Current biology : CB 13: 1116–1121.

Schaefer A, Sampath SC, Intrator A, Min A, Gertler TS, Surmeier DJ, Tarakhovsky A, Greengard P. 2009. Control of cognition and adaptive behavior by the GLP/G9a epigenetic suppressor complex. Neuron 64: 678–691.

Scheer S, Zaph C. 2017. The Lysine Methyltransferase G9a in Immune Cell Differentiation and Function. Front Immunol 8: 429.

Seisenberger S, Andrews S, Krueger F, Arand J, Walter J, Santos F, Popp C, Thienpont B, Dean W, Reik W. 2012. The dynamics of genome-wide DNA methylation reprogramming in mouse primordial germ cells. Mol Cell 48: 849–862.

Seki Y, Hayashi K, Itoh K, Mizugaki M, Saitou M, Matsui Y. 2005. Extensive and orderly reprogramming of genome-wide chromatin modifications associated with specification and early development of germ cells in mice. Dev Biol 278: 440–458.

Shinkai Y, Tachibana M. 2011. H3K9 methyltransferase G9a and the related molecule GLP. Genes Dev 25: 781–788.

Shirane K, Toh H, Kobayashi H, Miura F, Chiba H, Ito T, Kono T, Sasaki H. 2013. Mouse oocyte methylomes at base resolution reveal genome-wide accumulation of non-CpG methylation and role of DNA methyltransferases. PLoS Genet 9: e1003439.

Tachibana M, Matsumura Y, Fukuda M, Kimura H, Shinkai Y. 2008. G9a/GLP complexes independently mediate H3K9 and DNA methylation to silence transcription. EMBO J 27: 2681–2690.

Tachibana M, Sugimoto K, Fukushima T, Shinkai Y. 2001. Set domain-containing protein, G9a, is a novel lysine-preferring mammalian histone methyltransferase with hyperactivity and specific selectivity to lysines 9 and 27 of histone H3. J Biol Chem 276: 25309–25317.

Tachibana M, Sugimoto K, Nozaki M, Ueda J, Ohta T, Ohki M, Fukuda M, Takeda N, Niida H, Kato H et al. 2002. G9a histone methyltransferase plays a dominant role in euchromatic histone H3 lysine 9 methylation and is essential for early embryogenesis. Genes Dev 16: 1779–1791.

Tachibana M, Ueda J, Fukuda M, Takeda N, Ohta T, Iwanari H, Sakihama T, Kodama T, Hamakubo T, Shinkai Y. 2005. Histone methyltransferases G9a and GLP form heteromeric complexes and are both crucial for methylation of euchromatin at H3-K9. Genes Dev 19: 815-826.

Veselovska L, Smallwood SA, Saadeh H, Stewart KR, Krueger F, Maupetit-Mehouas S, Arnaud P, Tomizawa S, Andrews S, Kelsey G. 2015. Deep sequencing and de novo assembly of the mouse oocyte transcriptome define the contribution of transcription to the DNA methylation landscape. Genome biology 16: 209.

Xu Q, Xiang Y, Wang Q, Wang L, Brind’Amour J, Bogutz AB, Zhang Y, Zhang B, Yu G, Xia W et al. 2019. SETD2 regulates the maternal epigenome, genomic imprinting and embryonic development. Nat Genet 51: 844–856.

Zeng TB, Han L, Pierce N, Pfeifer GP, Szabo PE. 2019. EHMT2 and SETDB1 protect the maternal pronucleus from 5mC oxidation. Proc Natl Acad Sci U S A 116: 10834–10841.

Zuccotti M, Garagna S, Merico V, Monti M, Alberto Redi C. 2005. Chromatin organisation and nuclear architecture in growing mouse oocytes. Mol Cell Endocrinol 234: 11–17.

